# Detecting flying insects using mega-nets and meta-barcoding

**DOI:** 10.1101/2020.11.19.389742

**Authors:** Cecilie S. Svenningsen, Tobias Guldberg Frøslev, Jesper Bladt, Lene Bruhn Pedersen, Jonas Colling Larsen, Rasmus Ejrnæs, Camilla Fløjgaard, Anders Johannes Hansen, Jacob Heilmann-Clausen, Robert R. Dunn, Anders P. Tøttrup

## Abstract

Insect diversity and abundance are increasingly reported as being under pressure. However, monitoring insects across space and time is challenging, due to their vast taxonomic and functional diversity. This study demonstrates how nets mounted on rooftops of cars (car nets) and DNA metabarcoding can be applied to sample flying insect diversity across a large spatial scale within a limited time period.

During June 2018, 365 car net samples were collected by 151 volunteers during two daily time intervals on 218 routes in Denmark. Insect bulk samples were processed with a DNA metabarcoding protocol to estimate taxonomic composition, and the results were compared to known flying insect diversity and occurrence data.

We detected 15 out of 19 flying insect orders present in Denmark. Diptera. Psocoptera and Thysanoptera were overrepresented, while Hymenoptera, Coleoptera, Lepidoptera, Trichoptera, Odonata, Neuroptera and Plecoptera were underrepresented, compared to Danish estimates. We detected 319 species not known for Denmark. Our results indicate that this method can assess the flying insect fauna to a wide extent, but may be, like other methods, biased towards certain insect orders. Furthermore, car net sampling and DNA metabarcoding can update species records for Denmark while preserving specimens for natural history collections.

## Introduction

Recent studies have highlighted declines in the biomass and diversity of insects. These declines have come as a surprise, in part because of our poor understanding of spatial and temporal patterns in insect communities. One reason for this dearth is logistic; insects gathered via standardized sampling must be sorted manually. In biologically diverse regions, this is simply impossible, because most insect species have yet to be named (1) and thus the results of sorting one sample are very difficult to compare to the results of another sample.

One of the most commonly used approaches to sample flying insect diversity is the Malaise trap (2,3). Malaise traps sample flying insects at a point in space over a fixed time interval. These traps have the advantage of sampling many individual insects, but those species tend to come disproportionately from Diptera and Hymenoptera, two of the insect taxa most difficult to identify (4,5). In addition, malaise traps are difficult to use in landscapes in which most land is privately owned such as intensive agricultural fields and cities.

One complement to malaise trapping is car net sampling, which has the potential to sample in a way that integrates space (6,7). Rooftop or fender car nets have previously been used for targeted sampling of different types of flying insects, often with a focus on disease vectors (e.g. mosquitoes or black-flies (8–15), or beetles (16–18). Importantly, car nets can sample insects flying from both public and private lands, when the roads themselves are public.

### DNA metabarcoding for diversity assessments

Here we explore the addition of a novel innovation to car net sampling to overcome its chief existing barrier, sorting and identification. We seek to consider the potential of identifying the results of car net sampling through DNA metabarcoding techniques. Implementation of DNA metabarcoding techniques allows for fast and cost-effective processing of a large number of samples and can be used in monitoring programs for community assessment (19). With higher quality curated reference databases, large sample processing output and standardized monitoring schemes, DNA metabarcoding has the potential to become an applied method for insect diversity monitoring (20–22).

Our aim in this paper is to assess the car net sampling method and the application of DNA metabarcoding method to survey the proportional diversity of insects in Denmark. First, we test whether the proportion of taxonomic levels as well as within insect orders is equal to the proportions known for Denmark. Then, we examine whether we detect new species occurrences for the country and the region. Furthermore, we compute richness indices and estimates to assess species accumulation with increased car net sampling and identify core species communities in the samples. Finally, to put our results into perspective, we compare them with the overall yield from Malaise traps, which are often used to estimate flying insect diversity.

## Methods

Car net sampling was carried out by volunteers along 5 km manually designed routes during June 2018 in Denmark (Fig 1A). To cover activity periods and sites of as many species as possible, sampling was carried out within the time intervals 12:00-15:00 and 17:00-20:00, and routes were placed in forest, urban areas, farmland, wetland and grassland. Each route was driven from start to end point at a maximum speed of 50 km/h and then back to start again. Bulk insect samples were placed in 96 % ethanol and stored in a −20°C chest freezer prior to DNA extraction.

**Figure 1:**
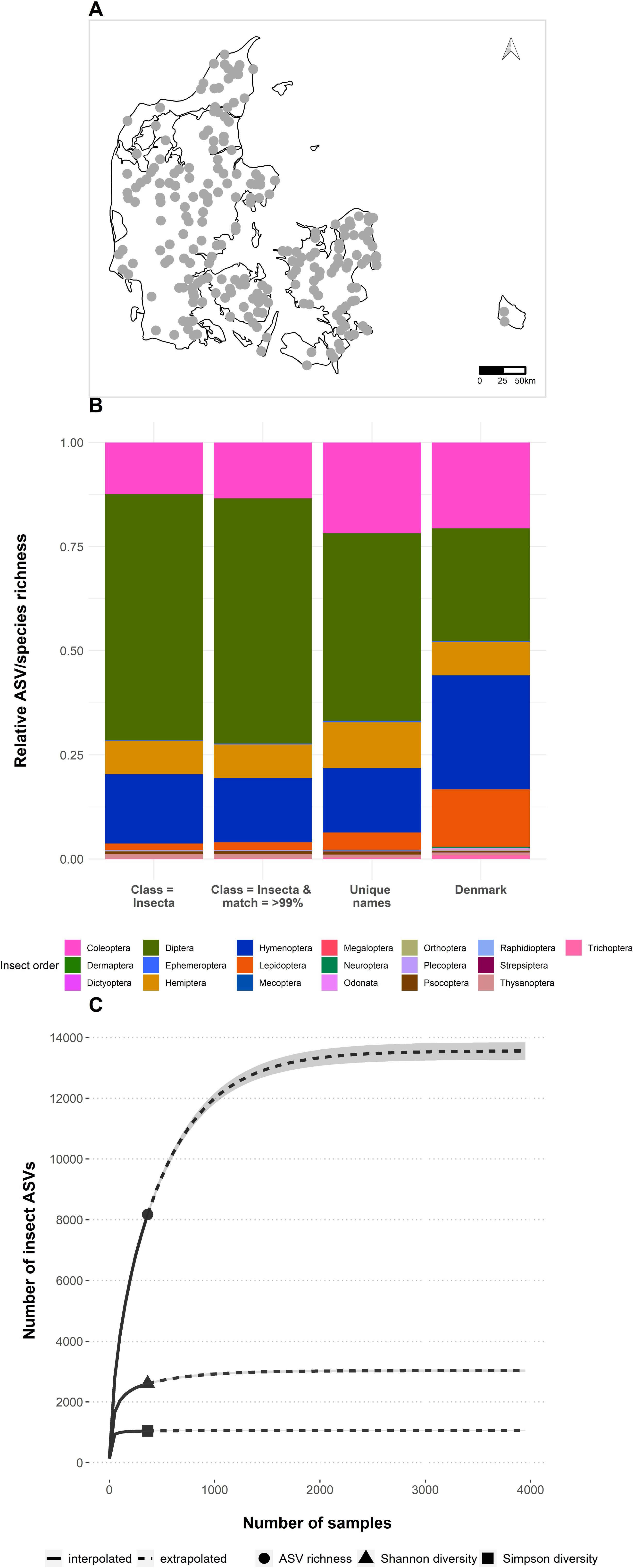
Sampling areas in Denmark (Northern Europe), relative species richness in car nets compared to the Danish species list and inter- and extrapolated species richness with increasing sample size. A) sampling areas in Denmark (Northern Europe) in June 2018. Flying insects were sampled along 5 km routes, driving back and forth to a total of 10 km. Points represent the route coordinate centroid. B) stacked barplots of relative species richness of insect orders for all car net samples compared to the relative species richness of insect orders known for Denmark, coloured by insect order. C) ASV accumulation curve by increasing sample size for observed richness, Shannon-Wiener diversity index and Simpson diversity index. The indices are included to account for community heterogeneity between. The grey fill area around the line illustrates the standard error.

DNA extraction, qPCR, PCR, library building and sequencing were carried out on 365 samples from 218 routes. The full protocol from sample processing to library build can be obtained here: dx.doi.org/10.17504/protocols.io.bmunk6ve, Data from the fwh primer pair are used in this study.

### Bioinformatics and statistical analysis

Samples from 14 libraries of 96 samples each were demultiplexed using cutadapt (version 1.11) (23). Amplicon sequence variants (ASVs) were identified and chimeras removed with DADA2 (24). Redundant sequences were removed with the LULU algorithm (25). The sequence ID tool from GBIF (https://www.gbif.org/tools/sequence-id) was used for taxonomic assignment. Further information on the bioinformatics analysis can be found in supplementary material.

All statistical analyses were carried out in RStudio (version 3.6.1). The ASV table was converted to presence/absence.

#### Comparison to the Danish species list and occurrence data

The proportion of insect ASVs in each insect order was compared to the Danish species list (www.allearter.dk), excluding the non-flying orders Phthiraptera, Siphonaptera, Zygentoma and Microcoryphia. We based the comparison on unique ASVs within each insect order filtered by match to class Insecta since we assumed unique reads to represent true diversity, but included comparisons to a ≥99% reference match and unique species names, thus eliminating sequence variants, for visualisation of differences in sequence and taxonomic diversity estimates.

We used a two proportions z-test, to test whether there were any differences in proportion of ASVs within insect order, family, genus and species, and amount of species detected within each insect order, between our results and the species in the Danish Species List.

We also investigated whether the car net samples contained species not registered in the public Danish species database or in the subset of data from the Global Biodiversity Information Facility from Denmark (26) and neighbouring countries Sweden, Norway and Germany (27). For reference database comparison, we based final taxonomic assignments on a ≥99% match across the entire query sequence and filtered based on unique species names. Species from our results without occurrences in Denmark or neighbouring countries were manually checked for occurrences in each country in Fauna Europea (28), and BOLD systems version 4 (29).

#### Species accumulation curve for car nets & core taxa

To investigate how many species the car net could potentially sample, we constructed ASV richness accumulation curves on row-sum frequencies of the species by sites presence-absence matrix with the package iNEXT (30,31). We visualised observed richness and accounted for community heterogeneity by including Shannon-Wiener and Simpson diversity indices. Additional richness estimates were calculated based on the transposed presence-absence matrix with the function poolaccum (vegan, version 2.5-6) (32) (Fig S2). Furthermore, we examined whether some ASVs were more abundant in our samples by extracting core ASVs present in 25% of all the samples using the RAM package (33).

## Results

From 365 car net samples, we identified 15 insect orders, 240 families, 1273 genera and 2114 species. This corresponds to equal proportions of flying insect orders, half of all families, a quarter of all genera and 11.3% of all species, on the complete Danish species list (Table S1 & Fig 1B).

Diptera was the most species rich group, followed by Hymenoptera, Coleoptera and Hemiptera, while Lepidoptera and Trichoptera were less represented in our samples compared to Danish estimates. The remaining insect orders were all represented by a proportion of <2% (Fig 1B). Equal proportions to the known diversity were obtained for Hemiptera, Orthoptera, Mecoptera, Dermaptera and Ephemeroptera (Table 1). The remaining insect orders were detected with 15% or lower proportion compared to Danish estimates. We did not detect any members of the flying insect orders Dictyoptera, Megaloptera, Raphidioptera and Strepsiptera in our samples.

**Table 1:** Results of the two proportion z-test results on proportion of ASVs/species in each insect order and the proportion of unique ASVs detected proportionally to known insect species for Denmark. Each insect order was tested against the known species number for Denmark within the insect order to assess whether our method detected equal or unequal proportions. Significant and unequal proportions are marked with bold.

| Insect order | Pearson's chi-squared ($\chi^2$) | P-value | lower CI | upper CI | Relative proportion of individual ASVs in car nets (%) | Relative proportion of individual species Denmark (%) | Proportion of unique ASVs detected in car nets compared to individual species in Denmark (%) |
| --- | --- | --- | --- | --- | --- | --- | --- |
| Diptera | 2509.2 | <0.001 | 0.31 | 0.33 | 59 | 27 | 94.9 |
| Hymenoptera | 361.46 | <0.001 | -0.12 | -0.1 | 17 | 27 | 26.4 |
| Coleoptera | 255.62 | <.0001 | -0.09 | -0.07 | 12 | 21 | 26.2 |
| Lepidoptera | 934.24 | <0.001 | -0.13 | -0.12 | 2 | 14 | 5 |
| Hemiptera | 0.03 | 0.86 | -0.01 | 0.01 | 8 | 8 | 43.9 |
| Trichoptera | 54.18 | <0.001 | -0.01 | -0.01 | 0.1 | 1 | 5.2 |
| Thysanoptera | 17.57 | <0.001 | 0.002 | 0.007 | 1.1 | 0.6 | 78.8 |
| Neuroptera | 21.17 | <0.001 | -0.004 | -0.002 | 0.02 | 0.3 | 3.2 |
| Psocoptera | 9.45 | 0.002 | 0.001 | 0.004 | 0.6 | 0.3 | 79 |
| Odonata | 8.96 | 0.003 | -0.003 | -0.001 | 0.1 | 0.3 | 15 |
| Orthoptera | 1.62 | 0.2 | -0.002 | 0.0002 | 0.1 | 0.2 | 26.3 |
| Ephemeroptera | 1.5 | 0.22 | -0.002 | 0.0003 | 0.1 | 0.2 | 27.9 |
| Plecoptera | 4.21 | 0.04 | -0.002 | 0.0002 | 0.04 | 0.1 | 12 |
| Dermaptera | 0.26* | 0.61 | -0.001 | 0.0002 | 0.01 | 0.03 | 16.7 |
| Mecoptera | 0.10* | 0.76 | -0.003 | 0.001 | 0.04 | 0.02 | 75 |

\* Pearson's chi-squared values were uncertain, due to the low number of taxa.

We found 319 species not registered in Denmark (Table S2). The majority of these (301 species) have occurrences in the neighbouring countries leaving 18 new species for the region (Table S3).

Observed ASV richness is extrapolated to increase from 8172 to 13154 unique ASVs (~38%) with further sampling (Fig 1C) with a sample coverage of 0.998 (95% CIs [0.998, 0.999]). The Shannon-Wiener and Simpson estimates of diversity, which to differing extents more highly weigh evenness, reached asymptotes within our sampling (Fig 1C). We found 15 ASVs present in 25% of all samples. These were assigned to eight well-known species and seven well-known genera (Table S4).

Sequence data results can be found in the supplementary material.

## Discussion

Using car nets and DNA metabarcoding, we detected insect orders in proportion to their diversity in the Danish species database (15 insect orders out of 19 known), including almost half of all flying insect families and 18 species with no occurrences from Denmark or neighbouring countries.

The insect orders Dictyoptera, Megaloptera, Raphidioptera and Strepsiptera were not detected in the car nets. However, species of Dictyoptera and Raphidioptera only take flight occasionally and usually stay close to vegetation (34,35), Megaloptera species tend to stay close to water bodies (35) and Strepsiptera females are endoparasites of insects and only the very short-lived males take flight (36). Since we sampled in the month of June 2018, we did not detect species that are present before and after June, and a longer sampling season will most likely increase the amount of species detected with our methodology.

For several years, the Swedish Malaise Trap Project (SMTP) has collected and morphologically identified flying insect diversity (3). Similar to our findings based on DNA, they found Diptera to be the most species rich group (75%), followed by Hymenoptera (15%), with less than 10% of the total catches belonging to the other insect orders and Acari. In our samples, we detected higher proportions of Hemiptera and Coleoptera, compared to SMTP. Although our results are based on one month’s sampling in 2018, it is strikingly similar in overall taxonomic composition to the multiyear national assessment for Sweden based on Malaise traps. Similarly to SMTP, we also find most of the new species detected belong to Diptera and Hymenoptera (38). However, our approach is much quicker and covers a much larger geographic area. Car nets can sample over 1000 individuals per sampling trip (~10-20 minutes) under favourable conditions which is similar to 24 hours of sampling with Malaise traps (4).

The DNA metabarcoding approach as employed here, or employed in concert with other sampling approaches, has definite advantages. However, given existing sequencing databases, it also has limits. First, several methodological steps in the laboratory inevitably introduce biases all the way from DNA extraction to sequencing, which have been well discussed elsewhere. Second, processed sequences need to be assigned to taxa (ASVs), the details of which depend upon algorithmic rules, which can influence the number of species identified as well as their boundaries. Finally, if ASVs are to be matched to morphologically named species (which they do not necessarily have to be for community and diversity assessments) DNA metabarcoding relies on updated reference databases and so is only as good as those databases. The databases for Denmark are relatively complete, but for many countries, particularly biodiverse countries, they are not.

Even in well-studied regions, reference databases differ among taxa. For example, Diptera and Hymenoptera have relatively fewer species references in BOLD compared to other insect orders, and small-bodied taxa are especially underrepresented (40). Since car nets catch large numbers of small Dipterans, DNA barcoding could in the future be used to generate references and fill the reference library gap for small-sized Diptera as well as other unknown taxa. The non-destructive DNA extraction method used in this study further has the advantage that single individuals can be isolated, morphologically identified and re-analysed to generate reference sequences.

One caveat with car nets as a monitoring tool is that certain weather conditions have to be present for sampling to be carried out, i.e. no rain and low wind speed, which makes it difficult to use in some areas that are prone to high wind speed, e.g. coastal areas. Another caveat is that since sampling is carried out using cars, roads have to be present and in good condition. Driving speed and the number of stops may have an impact on how many insects are sampled (41) and therefore requires routes to be designed with considerations of stops, turns and road conditions. For example, urban areas have a higher number of stop signs, road crossings etc. compared to rural areas, which in turn more often have gravel roads that requires the car to drive slower than on a paved road.

Nonetheless, for many uses, the limits and caveats associated with the combination of car nets and metabarcoding are outweighed by the benefits. By designing a simple, standardized citizen science project, we were able to sample at a large spatial scale within one month, with a response rate of more than 75% of the projected samples returned. As such, car net sampling with the help of citizen scientists could be a promising tool for monitoring flying insects across time and space.

## Conclusion

Car net sampling combined with DNA metabarcoding may be a useful tool for monitoring flying insect diversity and abundance across spatial and temporal scales. Furthermore, car nets can detect unregistered species, be applied to monitor selected taxa, e.g. mosquitoes, and have the potential application as a national monitoring method.

## Authors’ contributions

CSS, APT, RRD, RE, JHC, CF, JB designed the study, developed the sampling strategy, recruited citizen scientists, coordinated the sampling, carried out statistical analyses and drafted the manuscript. LBP carried out laboratory work and revised the manuscript. JCL designed and tested the car net, recruited citizen scientists and revised the manuscript. TGF, AJH contributed to the laboratory protocols, developed scripts for bioinformatics analysis and processing, and revised the manuscript. All authors gave final approval for publication and agreed to be held accountable for the work performed therein.

## Acknowledgements

We want to extend our sincerest gratitude to all the citizen scientists who contributed to the project. Furthermore, we wish to thank Thomas Pape for providing taxonomic expertise on the new species occurrences for Denmark and the neighbouring countries.

## Data Availability statement

Documentation of the analysis can be found on GitHub (https://github.com/CecSve/InsectMobile_CarNet). The datasets supporting the analysis in this article have been uploaded as part of the supplementary material.

