## Supplementary material for "Detecting flying insects using mega-nets and meta-barcoding": Table_S1

Table S1 The relative diversity of insect orders, families, genera and species in our samples compared to the Danish national species list.

| **Taxonomic level** | **Relative abundance of ASVs compared to Danish estimates (%)** | **Df** | **N** | *X*^2^ | **P-value** |
| --- | --- | --- | --- | --- | --- |
| Order | 79 | 1 | 19 | 2.52 | 0.11 |
| Family | 49.5 | 1 | 485 | 323.34 | <0.001 |
| Genus | 23.3 | 1 | 5467 | 6797.9 | <0.001 |
| Species | 11.3 | 1 | 18791 | 29974 | <0.001 |
