## Supplementary material for "Detecting flying insects using mega-nets and meta-barcoding": Table_S2

Table S2 List of species detected in the car net samples with no occurrences for Denmark on GBIF (September 23 2020) and the associated sequences used to assign taxonomy. The accepted name was extracted from GBIF back-bone taxonomy, however, the species may be present in Denmark under a synonym. The table is arranged alphabetically by order, family, genus and species.

| **ASVID** | **order** | **family** | **genus** | **species** | **sequence** |
| --- | --- | --- | --- | --- | --- |
| b7e02c3f301fe0f3dfd4fe8e5f68f2bd4be628f3 | Coleoptera | Ciidae | Ennearthron | Ennearthron cornutus | cctttcagctaatatttcacattctggatcatctgtagatatagcaattttcagattacatttagcaggagtatcttcaattcttggtgcagtaaattttatttcaacagtagttaatatacgtccaataggaataactttagatcgaataccattatttgtatgatcagttattattacagcaatcctattacttctttcatta |
| 74c056958dfe36dec4bbe5df3f5198c698e7b854 | Coleoptera | Coccinellidae | Cryptolaemus | Cryptolaemus montrouzieri | tttatcttctaatttagctcatagagggccttctgtagatttagctatttttagacttcatttagcaggtatttcatctattttaggagcaattaattttattactacaattttaaatatacgtccatatttaataacacttgaaaaaataccattatttgtatggtctgttttaattacagctgttcttcttcttctttcatta |
| c95623a83ac37938c2aeb560c27b8daf5b81fa68 | Coleoptera | Coccinellidae | Stethorus | Stethorus punctillum | tctttcaagtaatttagcgcatggtgggtcagcagtagatttatcaatttttagccttcatttagctgggatttcttctattttaggagcagttaattttattactaccattattaatatacgacctactggaataacattagataaaaccccattatttgtgtgatctgtgtttatcacagctattcttttacttctatcttta |
| ff5fe06c1a1cd738a31d80945c039f4a5f891125 | Coleoptera | Curculionidae | Bostrichus | Bostrichus cryptographus | tctagctgctaatattgctcatgaaggagcatcaattgatctagcaatttttagtttacatatatctggagcttcatcaatcctaggagcaattaactttatctcaacaattattaatatacacccaataggtataaaaccagaacaactatccctatttacatgagcagtaaaaattactgcaatcttacttctactttcactt |
| 8e75e8f343fbb50dfa14ea072b9cf617d6562c64 | Coleoptera | Curculionidae | Bostrichus | Bostrichus piceae | actagcatcaaacattgcccatgaaggagcatctgtagacttcgcaatttttagtttacatatagccggaatttcttcaatcttgggggcaataaattttatctctacaatcatcaacatacatcccagagggataaaaccagaacaactatcattattttcatgagctgtaaaaatcacggctatcctcctcttactatccctc |
| 78e42e538a0b672f36077aef851b98d9497f906a | Coleoptera | Curculionidae | Rhamphus | Rhamphus subaeneus | tttatcagctaatattgcccatgaaggaacttcagtagatttagctatttttaggctacatatagcaggagtctcatctatcttaggagcaataaactttatttcaaccattattaatataaaacctaaaaatataaaattagaacaaatatctctttttgtttgatctgtaaaaattacagctattttattattactttcacta |
| 33e581eda85be8f1e695940a42f26014281e4dfd | Coleoptera | Latridiidae | Enicmus | Enicmus brevicornis | tttatctagaaatattgctcatgggggatcatctgtagaccttgcaattttcagccttcacctagccgggatctcatctattctcggagccgtaaattttattactactgtaattaacatacgaccaacaggaataaaattagaacaattgcctttatttgtttgatcagtgaccctaacagcaattttactccttctctcatta |
| 24eeb820bf7f00237a7fbd7cb45ee3031a79463a | Coleoptera | Leiodidae | Liocyrtusa | Liocyrtusa minuta | cctctcttccaatattgctcatagaggatcatctgtagatttagctatttttagcttacatcttgcaggtatttcttctattctaggggcagtaaattttattacaaccgtaatcaatatgcgaccaacaggaatatcttttgataaaatacccttatttgtatgatcagttgttattactgcaattttgttgcttctatctctc |
| 564fa21ecb07fdb5b584f9c22e9ce9e8089c27d8 | Coleoptera | Staphylinidae | Amischa | Amischa forcipata | tctttcatctaatattgcccatggtggttcttctgttgatttagcaatttttagtcttcatttagctggaatttcgtcaattttaggagcagttaattttatttcaacagttattaatatacgatcaacaggaatttcatttgatcgaatgcctttatttgtttgatcagtagcaattacagcattacttcttcttctatcatta |
| b5b769dbd73f5bba215d96abbc8c3f6b42ae7e28 | Coleoptera | Staphylinidae | Anotylus | Anotylus clavatus | cctatcgtctaacattgctcatagaggatcttcagttgatttagctatttttagattacacttagctggaatctcttcaattttaggggcagttaactttatttcaactattatcaatatacgatcaattgggatatcctttgatcgaatacccctatttgtttggtctgttaatattacagctattttattacttctttcatta |
| 9256139116614301c29aba657d2b33fb5344affd | Coleoptera | Staphylinidae | Atheta | Atheta cribrata | cctttcttccaatattgcacatggaggatcttcagttgatctagcaatttttagacttcatctagcaggaatttcatcaattttaggagccgtaaatttcatttcaactattattaatatacgatccccaggtatttcatttgatcgaatacctctttttgtatgatcagtagcaattacagcattactactacttttatcctta |
| a97f115e333350cca3825842cf1073c289fa13ec | Coleoptera | Staphylinidae | Gabronthus | Gabronthus sulcifrons | cctatcctccaatattgcccatggaggagcatcagtagatctagcaattttcagccttcatttagccggaatttcatctattctaggagcagtaaattttattacgactgtaattaacatgcgatcaacaggaataacattcgaccgaatacccttattcgtttgatctgtagctattacagctctattacttctcctctctcta |
| d5965de7d1d944d5487df3f55c72e9be095c6abb | Coleoptera | Staphylinidae | Mocyta | Mocyta amplicollis | cctatcatccaatattgctcatggcggatcttcagttgatttggcaatttttagtctacatttagccggtatttcatctattttaggagcagtaaattttatctcaacagtaattaatatacgttcatcaggaattacatttgatcgtatacctttattcgtttgagcagtagctattacagctttacttttattactatcacta |
| 7cbcfd58fb3fcc0fdb2023785352351ae35a0683 | Coleoptera | Staphylinidae | Philonthus | Philonthus rotundicollis | actatcatctaatattgcccatggtggtgcctcagtagatctcgcaattttcagccttcatctagctggaatttcatcaatccttggggctgtaaattttattaccacagtaatcaacatacgatcgacaggaataaccttcgaccgaatacctttattcgtttgatctgtagctatcacagctttactgctactactatcactc |
| e57672d1054792f02888bba2c7c87d0dccda3c03 | Diptera | Agromyzidae | Amauromyza | Amauromyza karli | attatcttcaattattgctcatggaggagcttcagttgatttagctattttttctttacatttagccggagtttcttcaattttaggggcagtaaattttattactacaattattaatatacgaactactggaattacatttgatcgaatgcccctttttgtatgatctgtattaattacagctgtattattattactttcttta |
| 6a0dfc414f34d2a2da5b9c54ab31d05df85a9c98 | Diptera | Anthomyiidae | Botanophila | Botanophila relativa | tttatcttctaatattgctcatggtggagcttctgttgatttagctattttttctttacatttagcaggaatctcatctattttaggagctgtaaattttattacaactgtaattaatatacgatcaacaggaattacttttgatcgaataccattatttgtatgatctgtagtaattacagctttgttacttttattatcttta |
| c8b75d5591648e662c39c0ecc95164030c3c888d | Diptera | Anthomyiidae | Paradelia | Paradelia arctica | tttatcatctaatattgctcatggtggagcttctgttgatttagctattttctctcttcatttagcaggaatttcatcaattttaggagctgtaaattttattactactgtaattaatatacgatcaactggtattacttttgaccgaatacctttatttgtttgatcagtagtaattactgcattattacttcttttatcttta |
| 6fee4ed7c42d5232db5b708b54e0826480466800 | Diptera | Anthomyzidae | Anthomyza | Anthomyza pleuralis | actttcctctattattgcccatggaggagcatcagtagacttagctattttttcattacatttagctgggatttcctctattttaggggctgtaaatttcatcacaactgtaattaatatacgatcaactggaattacttttgaccgaatacctctatttgtatgatctgtagttattacagcttttttactcttactttcccta |
| a9679f683d76895bf2abd460b8ec4078bb98aaeb | Diptera | Asilidae | Eudioctria | Eudioctria propinqua | tctttcgtcaggaattgctcatagaggagcatctgttgatttagcaattttttcattacacttagctggaatttcctctattttaggagcagtaaattttattaccacaattattaatatacgatctacaggtattaaatttgaccgtatacctctatttgtatgatcagtaataattacagcaattttacttttattatcttta |
| 83bde943e03f5ff4c7d2acba8fb343d7ec9c1937 | Diptera | Asteiidae | Asteia | Asteia beata | actatcttctattattgcccatgggggggcttctgttgatttagctattttttcattacatttagcaggaatttcttcaattttaggagcagtaaattttattacaaccgttattaatatacgtgcaacaggaatttcatttgatcgaatacctttatttgtttgatcagtagtaattaccgcattattattactactatcattg |
| 48e1bf1d93c6487da5eee10984020ad78294b99a | Diptera | Canacidae | Pelomyia | Pelomyia occidentalis | gctttcttcaacaatcgctcatagaggagcatcagttgatttagcaattttttctctacatttagccggaatttcctctattttaggagctgtaaatttcattacaacagtgattaatatacgatcagtaggaattacttttgaccgaatacctctatttgtttgatcagtagtaattacagcccttttacttcttttatctctt |
| 0cac7f9a2a36c9f962899bee9f21a1e9b2955cd9 | Diptera | Cecidomyiidae | Aprionus | Aprionus subbetulae | ttgtcttcaaatattgctcattcaggaggagcagtagatttatctattttttctttacatttagcaggtatttcttcaattttaggggcaattaattttattactactataattaatatacaaacaaaaaaaattaaatttgaccaactatctttatttagttgatctgtagtcattactgcaatcttacttttgttatcttta |
| 2ed759f8537117d9fef1fbaee1ca865c05c56742 | Diptera | Cecidomyiidae | Campylomyza | Campylomyza aeratipennis | cctttcctcaaatttagctcattcaggaatttctgtagatttatcaatttttagcttacatttggcaggaatttcttctattttaggggcaattaatttcatttcaaccattttaaatataaaaattattaatataaaatttgattatttaatattatttatttgatctgttttaatcactgccacactacttttaatgtctttg |
| 2a94779687d3e44c261118bea1302e08724fe740 | Diptera | Cecidomyiidae | Leptosyna | Leptosyna nervosa | tctttcatcaattatttcacataatggagcatcagttgatttatctattttttctctccatttagcaggaatctcgtcaattcttggagctattaacttcattactacaataattaatatacgaattaaaataattaaatttgatcaaattccattatttgtttgatcaattattattactgctattcttttgctattgtctctc |
| 7b96e6736e1b1d7b46033db736f2ddd46629120d | Diptera | Cecidomyiidae | Parepidosis | Parepidosis argentifera | cttatcatctattattgctcacactggagctagagtagacctatctattttttctcttcatatagcaggaatttcttcaattttaggggccattaactttattactacaataattaacatacgaattaaaaatattaaatttgatcaaatccctttattttcatgatcagttattattacagccattttacttttactatcatta |
| c1184667fff302ec6d0a564090d940c6aed3da18 | Diptera | Cecidomyiidae | Peromyia | Peromyia cornuta | tctatcatctattattagacattctggttcatctgtagatttatctattttttcattacatatagccggaatttcttctattttaggggcaattaattttattactactttaattaatttacaaattatttctttaaaatttgatcaactttctttattttgttgatctgtttttattacagctattttattattattatcatta |
| d98f75c86ebdd505164b6cf2202e02887bfc0053 | Diptera | Cecidomyiidae | Peromyia | Peromyia impexa | gctatcctctgttattgctcattctggtacatcagtggatttttctattttttctttacatatagcaggaatttcttctattttaggggctattaattttatttcaacaatttttaatataaaattgaaatcaattaaattggatcaattaactttatttatttgatcagtaaaaattactgcaattttactattactttcactt |
| dc12593ce9f88159ae67e69763a6a294410b2140 | Diptera | Cecidomyiidae | Peromyia | Peromyia iuxtatruncata | cctatcttctactatagcacatctaggttcatctgtagatttatctattttttctttacatctagcaggaatttcatcaattttaggagcaatcaattttattactactataataaatatacaagtaaaaaatattaaatttgatcaattacctttatttgtctgatctgtaggtattactgcagtattattacttttatcctta |
| f002e14e6bd273770f0d502f37983f7c68526994 | Diptera | Cecidomyiidae | Peromyia | Peromyia ovalis | tttatcttcattattagcacattcagggccttcagtagatttatcaattttttctttacatttagctggaatttcttctattttaggagctattaatttcattactacaatattaaatatacaagtaataaatattaaatttgatcaattacctttatttacttgatctataattattactgcagtattattactactatcttta |
| 81cce6e1a19dde1b02abbce07506035ff969eb88 | Diptera | Cecidomyiidae | Peromyia | Peromyia photophila | tctttcctctattattgcgcattctggcaggtctgttgatttatcaattttttctttacacttagcaggaatttcatcaattctaggagcaattaattttatttcaactataataaatatacgagttttaaatattaagtacgaccaactacctctatttgtatgatcagttgtgattactgccattttactattattatcatta |
| 0811b8e5212e558db60b6e56b366fc7c71311add | Diptera | Cecidomyiidae | Peromyia | Peromyia ramosa | tctttcatctttattaagtcattctggatcttctgtagatttatcaattttttctcttcatttagctggaatttcttcaattttaggggctattaattttattactacaatattaaatatacaagtaaaaaaaattaattttgaccaattatcattatttacatgatctgtaattattactgctattttattattattatcttta |
| e6bd99d41ca1a0882dd15fae6d18329f3b3fabf2 | Diptera | Cecidomyiidae | Peromyia | Peromyia upupoides | cctttcatcagtaatttcgcatatcggggcttctgtagatttatctattttttctttacatttagcaggaatttcatcaattttaggagctattaactttatctctactataataaatatacaaattaactctattaaattagaacaattatccttatttatatgatctgttattattacagctatcctattacttctatcatta |
| 307897127bf24cb7cdd7b993db4d9167eef90a03 | Diptera | Cecidomyiidae | Porricondyla | Porricondyla colpodioides | ttatcatcaattttagctcataacggagcctcagttgatttatcaattttttctttacatatagccggaatttcctctatcttaggagctattaattttattactacaataattaatatacgtattaaaaatattaattttgatcaaattcctttattttcttgatcagtaattattactgctatcttattattactatctctg |
| 4b50a229725fa5c10c801bd87f326051bd1e17d6 | Diptera | Cecidomyiidae | Winnertzia | Winnertzia solidaginis | tctttcttcaattatttctcatacaggttcttctgtagatttatcaattttttcattacatctagccggtatttcttcaattctaggggcaattaattttattacaacatttattaatatacgaattaataatatcaaatttgatcaaatccctctatttacctgatcagtaataattacagcaattttacttcttctctcttta |
| 1ad3d64470756a03cc67a9b76f763a89f2ea6bcb | Diptera | Ceratopogonidae | Atrichopogon | Atrichopogon hirtidorsum | tttagccgctaatgtatcacatgccggagcgtccgtagatttagcaattttttctcttcatttagcgggaatttcatcaattttaggggctgtaaattttattactacaattattaatatacgatcaacaggaattacatttgaccgaatacctttatttgtctgatctgtatttattactgctattttactattattatcctta |
| d2e1b08f876c996db0c7a0cc8e5087dd6638d218 | Diptera | Ceratopogonidae | Atrichopogon | Atrichopogon oedemarum | tttagctgcaaatgtttcacatgcagggtcttcggttgatttagcaattttctctcttcatctagcaggaatttcttctattttaggtgcagtaaattttattactacaattattaatatgcgatcaaatggaattacattcgaccgaatacctttattcgtttgatctgtactaattacagctgttttactacttctttcctta |
| 25116669c9468029482c9e2112c32e7dbf0788ff | Diptera | Ceratopogonidae | Bezzia | Bezzia fuliginata | tttatcagctaatattgctcatgctggagcatctgttgatttagctattttttctttacatttagccggaatttcttcgattttaggagctgtaaattttattacaactattattaatatacgatcaaatggaattacttttgatcgaataccattatttgtatgatcagtattaattacagcaattttattattactatcttta |
| e2037be18e924a92f6f54299c17bb7ea3d07fae4 | Diptera | Ceratopogonidae | Brachypogon | Brachypogon bialoviesicus | cctttccgctaatatttctcatgcaggggcttcagtagatctagctattttttccttacatttagcaggtatttcatcaattttaggggcagtaaattttattactactattattaatatgcgctctaatggaattacatttgatcgaatgcccctatttgtttgatcagttttaattactgctatcctattactattatcttta |
| af2078e4f66c0fb9c0ba5a5c4a4880f4a9d4e729 | Diptera | Ceratopogonidae | Ceratopogon | Ceratopogon crassinervis | tttatctgccaatatctcccatgctggagcatctgtcgatcttgcaattttttcactacatctcgctggaatttcttccatcctaggagcagtaaacttcattactacgattattaatatgcggtctaatgggattacatttgatcgaatgcctctattcgtttggtccgttctaattaccgccatcttactacttttatccctc |
| 654bae4d6a5f2dfb0cf43d9a6e02731dd8acd074 | Diptera | Ceratopogonidae | Ceratopogon | Ceratopogon nitidulus | tttatcaactaacatctcccatgcgggggcctcagtagacttagctattttttctttacatttagctgggatttcctcaattttaggagctgtaaactttattactactattatcaatatacgttctaacggaattaccttcgaccgaatgcctctatttgtctgatcagtcttaattactgctattttattactattatcttta |
| d7e2ff627fd761f4d67243b0c9d7d85a9c316637 | Diptera | Ceratopogonidae | Ceratopogon | Ceratopogon perpusillus | ctctctgccaatatctcacatgcaggggcttcagttgatctagctattttttcccttcatttagccgggatttcttcgattctaggggcagtgaattttattaccactattattaatatacggtcaaatggaattacatttgaccgtatacctctttttgtgtgatccgtcttaattactgctatcctacttcttctatcttta |
| ec8f2e27960aed21f0e0409f1ee1120f9145e14d | Diptera | Ceratopogonidae | Ceratopogon | Ceratopogon sociabilis | cctttccgctaatatctcacatgccggagcctcagttgatctagctattttttctctccatttagccgggatttcttcaattttaggggcagtaaatttcattaccactatcattaacatacgatcaaacgggatcacattcgaccgcatacccctatttgtatgatctgttttaattactgcaatcttgcttcttttatccctc |
| 4c84161bd6e6c799fb99caa63ddaf728196d9813 | Diptera | Ceratopogonidae | Culicoides | Culicoides clintoni | tctttcagccaatgtttcacatgccggggcttctgtagatttagctattttttctttacatttagcgggaatctcttctattcttggagccgtaaattttattacaacaatcattaatatacgatctaatggaatttcatttgatcgaatacccctttttgtgtgatcagttttaattacagctattttattactactttcctta |
| 1ac9b97f4af2dda1ee1c8815896f768ec44c3920 | Diptera | Ceratopogonidae | Dasyhelea | Dasyhelea arenivaga | gttagcaagtaatcttgcccatagaggggcttcagttgacttagcaattttttctttacatttagctggaatctcttcaattttgggggcagttaattttattactactattattaatatacgatcaaatggtatttcatttgatcgaatgcctttatttgtgtgatctgttttaattacagcaattcttttacttttatcactt |
| a82e5ec935773d92078efaba7490559fa8441137 | Diptera | Ceratopogonidae | Dasyhelea | Dasyhelea aristolochiae | attagccgctaatatttctcatgcaggggcatctgttgacttagctattttttcattacatctagctggtatttcatcaattttaggagcagtaaattttattactactattattaatatacgttcaacaggtattacatttgaccggatacctctatttgtatgatctgttttaattacagctattttattattactttcacta |
| 7a0d4094801b8006ee24cd5aac8599f9ef950c63 | Diptera | Ceratopogonidae | Dasyhelea | Dasyhelea europaea | gttggcaaacaacatcgcccatagtggggcttctgttgatcttgcaattttttctcttcatctggccggtatctcatctatcctgggagcagtgaactttattacaactatcattaatatgcgttcaaaaggaatttcatttgatcgaatacccttatttgtttgatcagtccttattacagctattcttttacttctttcttta |
| bc89e9d8384ad3c23c518d126225c18cb6cbd998 | Diptera | Ceratopogonidae | Dasyhelea | Dasyhelea flavifrons | tctagctagtaacatagctcatggagggtcatcagttgatttagctattttttcccttcatttagctggaatctcctcaattctaggggctgtaaattttattactacaatcattaatatacgatcaaatggaattacatttgatcgtatacctttatttgtgtgatctgttcttattactgctgtattactattattatccttg |
| 852d16252258c0b6a6db1d2c8da6dc9d0b0045f4 | Diptera | Ceratopogonidae | Dasyhelea | Dasyhelea lucida | tttagctaataatatctctcatagagggtcatcggttgatttagctattttttctcttcatcttgcaggaatttcctctattttaggtgctgtaaattttattacaacaattatcaatatacgatctaaaggaatttcatttgatcgtatacctttatttgtatgatcagtcctaattactgcaattttattattgctttcatta |
| 29bf0b424e120ee83ee4ed069b53615e18a686e5 | Diptera | Ceratopogonidae | Dasyhelea | Dasyhelea turficola | tttagcaagtaatattgctcatagtgggtcatcagttgatttagcaattttttctcttcatttagcgggtatttcctctattttaggggcagtaaattttattactacaattattaatatacgctctaaaggaatctcatttgaccgaatacctttatttgtttgatctgtactaattacagccgtcttattattactttcttta |
| 6356a33f898ff10f8cb888ffe732dca0782dad6d | Diptera | Ceratopogonidae | Forcipomyia | Forcipomyia alacris | tttagccgctaatatctctcacacaggtgcttcagtagatttagctattttttcccttcatttagccgggatttcttcaattttaggtgctgtaaattttattaccactatcattaatatacgatcagtgggaattacatttgatcgaatacctttatttgtttgatctgttttaattacagctattttactattattgtcttta |
| 8856622aaa5c37f2c7d32d7f6a5dfe12273421fd | Diptera | Ceratopogonidae | Forcipomyia | Forcipomyia chaetoptera | tttagcagccaatgtttctcacgcgggatcttcagtagatttagctattttttctttacatttagcaggaatttcttctattttgggggctgtaaattttattacaactgtaattaatatacgatccgcaggaattacatttgatcgaatgcctttatttgtatgatctgttttaattactgcagttttacttttattatctctc |
| 5064fc06229da7755fa558f7ef0cc6cdf701392c | Diptera | Ceratopogonidae | Forcipomyia | Forcipomyia knockensis | tttagccgctaatatttctcacgcgggatcttcagttgatttagctattttttcactacatttagccgggatttcttcaattttaggggcagtaaattttattacaacaattattaatatacggtcctcaggaattacatttgatcgaatacctttatttgtttgatcagttttaattactgccattttattacttttatcttta |
| a6c38ff6af6c98ccb99d623e1198f69366c1fc7d | Diptera | Ceratopogonidae | Forcipomyia | Forcipomyia murina | tcttgcagctaacatctctcatgccgggtcttcggttgatttagctattttttcacttcatttggcgggtatttcatcaattttaggggcagtaaattttattactacaattattaatatacgttcaattggaattacatttgatcgaatacctttatttgtgtggtcagttttaattacagctattcttttacttttatcctta |
| 820125a9fa95c64e3440b50af6b2280169df44a3 | Diptera | Ceratopogonidae | Forcipomyia | Forcipomyia pulchrithorax | ttagccgccaatatttctcacgccggatcatccgtcgacttcgccattttctctctccatttagcaggaatttcatctattttaggagcggttaatttcattacaacaattattaatatgcgatcaaatggaattacatttgaccgtatgcctctttttgtatgatctgtattaattaccgcagtgttattacttttatcttta |
| d4c1932282eb4685cc7097c3dfcbb760795c3803 | Diptera | Ceratopogonidae | Forcipomyia | Forcipomyia squamigera | tctagccgccaatatttctcatgctgggtcatcagtagacttcgcaattttttctctccatttagctgggatttcttcaattctaggggcagtaaattttattactacaattattaatatacgatctaatgggattacctttgaccggatgcccctttttgtgtgatctgtactaattacggctattttacttcttctttcccta |
| e78bd27c7e29d6ca80009e18d76015ecfe0f9ecd | Diptera | Ceratopogonidae | Forcipomyia | Forcipomyia tenuis | attagctgccaatatctctcatgcaggctcttctgttgattttgctattttttccctccatttagccggtatttcctcaattttgggggccgttaattttattacaactattattaacatacgatcaaatggaattactttcgatcgcataccattatttgtatgatcagtcctaattactgcaattcttcttcttttatcttta |
| 488558a680a81c633217c9a9fc3d970df18bf90a | Diptera | Ceratopogonidae | Forcipomyia | Forcipomyia titillans | attagctgcaaatgtatcacatgccggggcctccgtagatttagcaattttttctttacatttagcaggtatttcttctattttaggggctgtaaattttattacaacaattattaatatacgatcagtggggattacatttgatcggattcctttatttgtttgatcggttttaattactgcaatcttactacttttatcatta |
| b3997e5a60ce3b97eddb93a45cd27ea012e3aed7 | Diptera | Ceratopogonidae | Forcipomyia | Forcipomyia velox | tttagcttctaatatttctcatgcaggtgcttcagtagatttggcaattttttctttacatttagccggaatttcttcaattttaggggctgtaaattttattaccacagtaattaatatgcgagctaacggaatttcttttgatcgaatacctttatttgtttgatcagtattaattactgcaattcttttattattatcttta |
| 0217d085689bcb62a7403d102999a9c192052896 | Diptera | Ceratopogonidae | Stilobezzia | Stilobezzia gracilis | tctttcagcaaatgtgtcacatgccggcgcatctgtagacctagcaattttttcccttcacttagccgggatttcttccattctaggagctgtaaactttattacaacgattattaatatacgatctaatggaattacatttgaccgaatacccctatttgtatgatctgtccttattacggctattctattattgctatcatta |
| c0578441dee498a8816e9d96cd5776b3914b1778 | Diptera | Ceratopogonidae | Stilobezzia | Stilobezzia ochracea | cctttcagctaatgtatcccatgcaggagcatctgtagatttagctattttttcccttcatttagcaggaatttcgtcaattttaggagcagtaaattttattacaactattattaacatgcgatctaatggaattacttttgatcgtatgccgctatttgtctgatctgttcttattacagctatcttattactactatctttg |
| b9781a8c143f0037b13d14ea59fec7a5f876b955 | Diptera | Chironomidae | Bryophaenocladius | Bryophaenocladius nitidicollis | tctctcttctaatattgcccacgcaggaagatctgttgacttagcaattttttcacttcatatagcaggaatttcttcaattttaggcgctattaattttattactacaattattaatatacgacctaagggaatatccatggaacaaatacctttatttgtttgatctgtttttattacagctattctccttttactttctctc |
| 383e7c69384f72c4114ab157c19cadba5a00326f | Diptera | Chironomidae | Bryophaenocladius | Bryophaenocladius subparallelus | tttatcatcaggaatcgctcatactgggggatctgtagatttagctattttttctttacacttagctgggatttcttctattttaggagcagtaaatttcattaccacagtaattaatatacgatcagaaggtattacatttgatcgaatacccttatttgtttgatctgtagtaattactgcaattttattattactttcttta |
| b0bfa92d3cba95aaba665ed36f38cc9bbbcbd545 | Diptera | Chironomidae | Chaetocladius | Chaetocladius melaleucus | tttatcctcaggaattgctcacgcaggagcatcagttgacctagctattttttctttacatttagcaggtatttcttctattttaggagcagtaaattttattactacagtgattaatatacggtccgaaggaattacttttgatcgaatacctttatttgtttgatctgtagtaattactgctattttactacttttatcttta |
| aebe5f238592870c883df991fa166462d4f81a29 | Diptera | Chironomidae | Chironomus | Chironomus atripes | tctttcttcaggaattgctcatactggaggttctgtagatttagcaatcttttcgttacatttagcgggtatctcatctattttaggagcagtaaattttattacaactgttattaatatacgatcaaatggaattacattagaccgaatacctttatttgtttggtcagtaattatcacagctattttattacttttatcctta |
| bfe939b388b6c6d09a32347a3d6df3749a7bd307 | Diptera | Chironomidae | Chironomus | Chironomus intersectus | tttatcttcaggaattgcccatgcaggagcatctgttgatttagctattttttctttacatttagcaggaatttctagtattttaggagctgtaaattttattacaacagtaattaatatacgatcagaaggaattactttagatcgaatacccttgtttgtttgatctgttattattacagctattttattacttctatcatta |
| 7e86319c42c60e82bcbcf14b2d55de5c0abf2e53 | Diptera | Chironomidae | Chironomus | Chironomus melanescens | cctgtcttctgctattgcccacagaggtgcctcagttgatttagcaatcttttctttacatttagcgggaatttcatctattcttggatcagtaaatttcattacaactgttattaatatacgggctaatggaattaccctagatcgaataccattatttgtttgatcagttgtaattactacagttcttttacttctttcactt |
| 5583bc90b55a5a218a25731bc5bb848873426fbc | Diptera | Chironomidae | Chironomus | Chironomus pseudomendax | tctttcttcagcaattgcccacagaggggcctcagtagatttagctattttttcattacatttagcaggtatctcatcaattcttggttctgtaaattttattacaactgtaattaatatgcgggctaatggtattaccttagaccgaatacctttatttgtttggtccgttgtaattacaacagtccttcttttactttcttta |
| c3fc6102b1400384dfb712357815e534c61c566a | Diptera | Chironomidae | Chironomus | Chironomus pseudothummi | tctttcatctgctattgcccatagaggagcatcagtagatttagctattttttctcttcatctcgctggagtttcatccattctaggttctgtaaattttattactacagttattaatatacgagcaaatggaattacactagatcgaatgcctttatttgtttgatcagtagttattactactgtattactactattatcttta |
| f2a370c3e307e7f5d5e9223d488ba1cadfba1dab | Diptera | Chironomidae | Chironomus | Chironomus punctatus | tttatcatccggaattgcccatgcaggtgcttctgtcgacttagcaattttttctctacacctagctggtatttcctctattttaggagcagttaattttattaccactgttattaatatacgttctaatggaattactttagaccgtataccattatttgtatgatccgttgtaattactgcaattttacttttgttgtcttta |
| 0ffea92897ddad93787566fcd6d7dbd1bbd9cacd | Diptera | Chironomidae | Cladotanytarsus | Cladotanytarsus gedanicus | actatcctcaaatattgcacatagaggagcttccgttgatttagctattttctcccttcatttagctggaatttcttctattttagggtcagtaaattttattaccacagcaattaatatacgaagaaacggaattactttagatcgaatgcccttattcgtttgatcagtagtgattacgacaattcttcttttattatccttg |
| ae1c79d7a99c3d237ae2dfd5f45fb1f282f3699e | Diptera | Chironomidae | Cladotanytarsus | Cladotanytarsus pallidus | tttatcctcaaatattgcacatagaggagcttctgttgatttagctattttttcccttcacttagctggaatctcatccattttagggtctgtaaacttcattactacagcaattaacatacgaagaaatggaattacactagaccgaatacctttattcgtctgatctgtaattattaccactattctacttttattgtcgttg |
| 9432fa33814d275a751b108b052cd18854af11dc | Diptera | Chironomidae | Cladotanytarsus | Cladotanytarsus wexionensis | tttatcttcaaatatcgcacatagaggggcttctgttgatttagcaattttctctcttcatttagctggaatctcttctattttaggttcagtaaattttattactacagcaattaatatacgtagaaatggtattaccctagatcggatacccctatttgtctgatctgtcattattaccactatcctacttttattgtcttta |
| a722ead9d30a22998636c564f099cd5e85c7e452 | Diptera | Chironomidae | Corynoneura | Corynoneura arctica | tctatcagcaaatattgcccatgcaggagcttcagttgacttagcaattttttctcttcatttagcagggatttcttcaatcttaggagcagtaaattttattacaacagtaattaatatacgttcagaaggaatttctttagatcgaatacctctttttgtgtgatccgtcgtgattacagcagtcttacttttactatcttta |
| 54449f2f029c676a4e9320072e2cdd240b304ffb | Diptera | Chironomidae | Corynoneura | Corynoneura gratias | tttatcatctaacattgctcatgctggcgcttcggttgatttagcaattttttctttgcatttagcagggatttcttcaattttaggagcagtaaattttatcacaacagtaattaatatacgatcagaaggaatttctttagatcgaataccactttttgtttgatcagtagtaattacagccgttctattacttctttcttta |
| ed2fd860bd028c9a43aedb646bf149daf075f8cb | Diptera | Chironomidae | Cricotopus | Cricotopus laricomalis | tttatcttcaggtattgcccatgcaggtgcatccgtagatttagctattttttcattacatttagcaggaatttctagaatcctaggagccgtaaattttattactactgtaattaatatacgttcagaaggaattactctagaccgaatacccttatttgtttgatctgttattattacagctattttactacttttatcatta |
| 56949660fd47daf56322231b5561dd416e7bfdfb | Diptera | Chironomidae | Cricotopus | Cricotopus reversus | tctttcatctggaattgcccatgctggagcttctgtagatttagctattttttctcttcatttagcgggaatttctagaattttaggagctgtaaattttatcacaactgttatcaatatacgatctgaaggaatcacattagaccgaatacctctatttgtttgatcagttattattacagccattttacttttactttcatta |
| 51b1a6eb870bb2a4370b0a9c4afadd6df35b29f6 | Diptera | Chironomidae | Cricotopus | Cricotopus rufiventris | tttatcttcaggaattgcacatgctggagcttctgttgacttagctattttctctttacatttagcaggaatttcctcaattttaggagcagtaaattttatcacaacagtaattaatatacgatcagaaggaattactctagaccgaatacccttatttgtttgatccgttgtaatcactgctattcttcttttattatctcta |
| 50629aedafdc0fdf8c83c66a9e601a8ae432f813 | Diptera | Chironomidae | Glyptotendipes | Glyptotendipes signatus | cttatctgccgctattgcacatagaggagcttcagtagatttagctattttttctttacatttagcaggtgtatcctcaattttaggttcagtaaattttattacaacagttattaatatgcgagctaatggtattacactagaccgaatacccttatttgtttgatctgtagtaattactacagtactattacttctttcttta |
| 63072c59fbeb01eb1c316d5b7572f5c89d27dd09 | Diptera | Chironomidae | Limnophyes | Limnophyes asquamatus | tctttcagctagtattactcatgctggagcctcggttgatttaactattttttctttacatttagcaggaatttcttctattttaggggcagttaattttatcacaactgtaattaatatgcgatcagaaggaattacttttgatcgaatacctctatttgtgtgatcagtattaattacagctattttattgcttatttcatta |
| 0a1f44f2bbb19a1aeca94a34348a003f0e294cdc | Diptera | Chironomidae | Limnophyes | Limnophyes natalensis | cctttcttcaagaatctctcatgcaggggcttctgtagatttagctattttttcattacatctcgccgggatttcctcaattctaggagcagttaattttattaccactgtaattaatatgcggtcagaaggaattactttcgatcgaatacctctttttgtttggtctgttttaattacagctattctattacttctctcttta |
| 8890a3c4e6dbe3d6a98a1fc42a5abc3c98d349b7 | Diptera | Chironomidae | Limnophyes | Limnophyes prolongatus | tctttcttctagaatttcacatgctggagcctcagttgacttagctattttttctttacatttagcgggtatttcttcaattttaggagctgtaaattttattactacagtaattaacatacgatctgaaggaattacttttgatcgaatacctctatttgtttgatcagttttaattacagctattcttcttcttctttcatta |
| 720cb22ab5a7fc035bba766fc4bcccb21a058444 | Diptera | Chironomidae | Metriocnemus | Metriocnemus atriclava | actatcttcaagaattgctcatgctggcgcttctgttgacttagcgattttttcccttcacttagctggaatttcttcaattcttggagcagtaaattttattactacagtaattaatatacggtcagaaggaatttccctcgatcgaatacctttgtttgtatgatcagtagtaattaccgccgttcttttattactttcttta |
| b2f445612d3ecb6aecada9a1b8ddfe4115d4f24f | Diptera | Chironomidae | Micropsectra | Micropsectra pallidula | tttatcttcaagaattgctcatagaggagcttctgtagatttagctattttttctcttcatctggctggaatttcttctattcttggttctgtaaattttattactacagctattaatatacgttcaaatggaattacattggatcgaatacctttatttgtttgatcggttattattactacagttttattacttttatcattg |
| 308eaa422a40f4737f453547e7106cc9ea963ea5 | Diptera | Chironomidae | Parachironomus | Parachironomus parilis | tctttcttctgcaattgcacatagaggagcttcagttgatctagcgattttttctcttcatttagctggagtatcttcaattttaggttcagtaaattttattactacagttattaatatacgagcaaatggaattacattagatcgaatacctctttttgtttggtctgttgtaattacaactgtcttacttcttttatctctt |
| be2ad9e229baad587bd9f9439a781834a89616cf | Diptera | Chironomidae | Paracladius | Paracladius quadrinodosus | tctatcttctgggattgctcatgctggagcttctgttgatttagctatcttttctttacatttagcaggaatttcttcaattttaggagctgtaaattttattacaacagtaattaatatacgatctgaaggtattactttagaccgaataccgttatttgtttgatctgttgttattacagctattcttttattattatcttta |
| 440ec8d7f3fbbeb4f1040b2b0478851a5b562266 | Diptera | Chironomidae | Paralimnophyes | Paralimnophyes hydrophilus | tctttcttctagaattgctcatgcaggagcttctgtcgatttagcaattttttctcttcatttagcaggtatctcatctattttaggtgccgtgaattttattactacagtagttaacatacgttcagaaggaatttcttttgatcgaatgcctttatttgtatgatctgttttaattacagctattttacttcttctttctcta |
| aabab5c7306dc70e14181b4bf5915f556b6843ce | Diptera | Chironomidae | Paratanytarsus | Paratanytarsus laetipes | tttatcttctagaatcgcccatagaggagcttcagttgacttagcaattttttcattacatctagcaggtatttcttctattttaggttctgtaaattttattacaacagcaattaatatacgttcaaatggtattacattagatcgtatacctttatttgtctgatctgtaattattacaacaattctattacttctttctctt |
| 1c541820ae8688f6eec21feb3f8c63c56296340b | Diptera | Chironomidae | Polypedilum | Polypedilum uncinatum | tctttcagctagtattgctcatagaggagcttctgttgatttagctattttttctcttcatcttgcgggagtatcctctattctaggatccgttaattttattacaacagtaattaatatacgttcaaaaggaattactctagaccgaatacctttatttgtgtgatctattgtaattacaacagttcttttacttttatcttta |
| e201a7aa261ca7ed2ae067ccbda18d011433f54e | Diptera | Chironomidae | Procladius | Procladius sagittalis | tctagcttcaggaattgctcatgcaggtgcttctgttgatttagcaattttttctcttcatttagctggagtttcttctattttaggtgccgtaaattttattactacagtaattaatatacgatctaacggaatcactttagaccgaatacctttatttgtatgatcagttgttattacagcagtattattacttttatcttta |
| e44fa0838e618c3a2b8cfe118c8a0ebdfdf65f74 | Diptera | Chironomidae | Psectrocladius | Psectrocladius conjungens | tttatcctcaggtattgctcatgcagggggctctgtagatttagcgattttttctttacacttggctggaatttcctctattttaggggctgtaaattttatcactacagtaattaacatacgatcaaatggtattactttagaccgaatacctctatttgtgtgatctgttgtaattacagctattctattacttctttcctta |
| 386eb6410370e0afda39772351c25182f7a3d22a | Diptera | Chironomidae | Pseudosmittia | Pseudosmittia danconai | actatcatctagaattgctcatagcgggggatctgttgatttagcaattttttctttacacttagcaggaatttcctcaattttaggagctgtaaattttattactacaattattaatatacgatcagaaggaattacttttgatcgaatacctttatttgtttgatcagtttttattactgcgatcttattattactttcttta |
| b573919b11223a3532f75a81871b49fa1a309000 | Diptera | Chironomidae | Pseudosmittia | Pseudosmittia jemtlandica | tctttcggcaagaattgctcatgctggagcttctgtagatttagctattttttctttacacttagctggaatttcttctattttaggagctgtaaactttattactacagtaattaatatacgatcaaatgggttaacttttgatcggatgcctctatttgtctgatcagtattaattacagcaattttacttcttctttcttta |
| 24220bbaba5b998f18c514f53c682908b29c48ee | Diptera | Chironomidae | Pseudosmittia | Pseudosmittia trilobata | tttatcttctagaattgctcacgctgggggatctgtagatttagctattttttctcttcatttagcaggaatctcttctattctaggagcagttaattttattacaacagtaatcaatatacgatcagaaggaattacatttgaccgaatacctctatttgtttggtctgttttaattacagcagttttattattactttcttta |
| 1e6dac666d5acfd0c91ef7deb2d8f5d116ae7671 | Diptera | Chironomidae | Smittia | Smittia albipennis | cttatctgcaggaattgctcatgcaggaggttctgttgatttagcaatcttctctttacacttagcaggaatttcctcaattctaggggcagtaaattttattacaactatcattaatatacgttctacgggaatttctttcgaccgcatacctctatttgtttggtctgtcctaattaccgctgttttactgcttctttctctt |
| 9d65078b462cfc904a725b0aa2c287a4c2524ca0 | Diptera | Chironomidae | Tanytarsus | Tanytarsus dissimilis | attatcttcaagaattgctcatagaggggcctcagttgatttagcaattttctctcttcacttagcaggaatttcttctattttaggttcagtaaattttattacaacagctattaatatacgatcaaatggaattactttagatcggatacctttatttgtttgatctgtaattattactactattttacttcttttatctctt |
| d39626e7601bcb3c1224f0d93931b7797c1539b1 | Diptera | Chironomidae | Tanytarsus | Tanytarsus excavatus | cctttctgctaatattgcccatagaggagcatccgtagatcttgctattttttctttacatttagcaggaatttcttcaattttaggctcagtaaattttattactacagctattaatatacgagctaatggaatcactctagatcgaatacctttatttgtctgatctgtaattattacaacaattttacttcttctttctcta |
| ffaeeb8d13d3a04ee9220505ed87502b5f8f291b | Diptera | Chironomidae | Tanytarsus | Tanytarsus heusdensis | gttatccgctagaattgcacacagaggtgcttctgtagacctagctattttttctctacatttagccggaatttcttcaattttaggatcggtaaattttattacaactgcaattaacatacgatcaaatggaattactcttgaccgtatacctttatttgtatgatccgtagtaattacaacaatcttactccttctctcccta |
| c9bc40e7c51e53ccc613609fd46cd0214098f4b1 | Diptera | Chironomidae | Tanytarsus | Tanytarsus lestagei | tttatcagctagtattgcccatagaggagcatctgttgatttagctattttttccctccatttagcaggaatctcatcaattttaggatcagtaaattttattactacagctattaatatacgggctaatggaatcactttagatcgtatacctttatttgtttgatctgttgttattactacaattcttcttcttttatcttta |
| 5f2a7aecea051d098db922606c5602ce51292bc0 | Diptera | Chironomidae | Tanytarsus | Tanytarsus volgensis | tatcatctagaattgctcacagaggagcatccgttgatctcgccatcttttcacttcatttagcaggaatctcatcaattttaggttccgtaaattttattacaactgcaatcaatatacgatcaaatgggatcacactggatcgaatacctctgtttgtgtgatccgtagtaatcaccactattttactccttctttctctc |
| 9b4e66590336e836e402c69b396f0ed02e8d04f2 | Diptera | Chironomidae | Thienemannia | Thienemannia gracei | tctctcgtctagaattgctcatgcgggtgcctcagtagatttagcaattttttcgttacatttagccggtatttcttccattttaggggcagtaaattttattacgacagtaattaatatgcgatctgaaggaatttcttttgatcgtatacctttattcgtttgatcagttgttattactgcgattttattattattatctctc |
| 4f605b35be777e14c1ab1436e833432c139e1c26 | Diptera | Chironomidae | Thienemanniella | Thienemanniella caspersi | tctttcttctaatattgcccatgccggagcttcagtagatttagcaattttttctcttcatttagcaggaatttcttcaattttaggagctgtaaattttatcactacagtaattaatatacgatctgagggaattagtttagaccgaatacctctatttgtttgatcagtaattattacagcaattttattattactatcttta |
| 6853cf6be6710d00aac114aabc0cbf6dbd8791e1 | Diptera | Chironomidae | Thienemanniella | Thienemanniella obscura | tttatcttcaaatattgctcatgcaggagcttcagtagacttagctattttttctctccatttagcgggtatttcgtcaattttaggagcagtaaattttattacaactgttattaacatacgatctgaaggaatcacactagatcgaatacccttatttgtatgatcagtcattattactgctattcttcttcttctttcttta |
| 7f5b228c6cdef128fb26c956274fc403e0b15288 | Diptera | Chloropidae | Chlorops | Chlorops nitidissima | tttatcttctattattgcacatggaggggcttcagttgatttagctattttttctcttcatttagctggagtatcatcaattttaggagcagtaaattttattactacagtaattaatatacgatcaacaggaattacatttgatcgaatgccattatttgtgtgatcagtagttattacagctctattattattactatcttta |
| d493e3502972c42b66b257dd0ec2725bf14afdd1 | Diptera | Chloropidae | Elachiptera | Elachiptera decipiens | tctatcttcaattattgctcatggaggtgcatcagttgacttagctattttttcattacatttagctggtgtatcatcaattttaggagcagtaaattttattactacagtaattaatatacgatcaacaggaattacatttgatcgaatacctttatttgtatgatcagttgtaattacagcattattattattattatcatta |
| 65e21a602fc96571881cafbd2fd870b0e003fbf0 | Diptera | Chloropidae | Oscinisoma | Oscinisoma alienum | tctttcttcaattattgctcacggaggagcttcagttgatttagcaattttttctttacacttagctggagtttcatcaattttaggggcggttaactttattactacagtaattaatatacgatcaactggaattacatttgatcgaatacctttatttgtttgatcagttgtaattactgccctattattactactatcttta |
| f3033f4a8a6fd9220c6cc318cf1521ac45ac8ca8 | Diptera | Chloropidae | Rhopalopterum | Rhopalopterum carbonarium | actatcttctattattgctcatggaggagcatcagttgatttagctattttttctcttcatttagctggagtatcttctattttaggagctgttaattttattacaacagtaattaatatacgatcaacaggaattacatttgatcgaataccattatttgtatgatcagtagttattactgctttattattattattatcttta |
| f3781052c6b6dc78a43edb98c70e5d2f37df335b | Diptera | Chloropidae | Thaumatomyia | Thaumatomyia pulla | cctttcttctattattgcccatggaggagcttctgtagatttagcaattttttccttacatttagcaggaatttcttcaattttaggagcagtaaattttattactactgtaattaatatacgatctacaggaattacctttgatcgaataccattatttgtttgatcagttgtaattactgctttattattacttttatcttta |
| 03e5dbb58f5f2143218b3a1814bb14d2ff13f0f1 | Diptera | Chyromyidae | Gymnochiromyia | Gymnochiromyia concolor | tctttctagaggaattgctcatggaggtgcatctgttgatttagctattttttcattacatttagctggaatttcttcaattttaggagccgtaaattttattactacagtaataaacatacgatcaacaggaattacatttgatcgaatacctttatttgtttgatcagtagtaattaccgctttattattactactttcatta |
| 9fbd0ec707c6b4cbc0a7693feaa470c8c58ed5f9 | Diptera | Cypselosomatidae | Pseudopomyza | Pseudopomyza atrimana | tttatcttcagttattgctcatggaggagcttcagttgatttagctattttttctctacacttggcaggagcttcttctattttaggggcagtaaattttattacaacagtaattaatatacgatctactggaattacactagatcgtatacctttatttgtgtgagctgtagtaattacagcttttttattacttttatctcta |
| 91034c07c6982ab28ea57dc88a34dc68da2e7f2d | Diptera | Dolichopodidae | Campsicnemus | Campsicnemus alexanderi | tttatctgcaggaattgctcacggaggtgcttctgttgatttagctattttctctcttcatttagccggaatttcttctattctaggagctgtaaattttattacaacagtaattaatatacgatcaacaggaattacttttgatcgtataccattatttgtatgatctgttgttattactgctattttattattactttcatta |
| c9ede0d235793ac474f8961c19ea85455e3a41d5 | Diptera | Dolichopodidae | Dolichopus | Dolichopus celeripes | cttatcctcaggaattgctcatggaggagcatctgtagatttagcaattttttcattacatttagcaggtatttcctcaattttaggggcagtaaattttattacaacagttattaatatacgatctacaggaattacatttgatcgaatacccctttttgtatgatcagtagtaattactgctattttattgttattatccctc |
| 0c9cd8046ca1be1c60657125d1eb708bdecc3a63 | Diptera | Dolichopodidae | Medetera | Medetera pseudoapicalis | tctttctagtggtattgcccatgggggagcttcagtagacctagcaattttctccctccacttagcaggaatttcctctattctaggagccgtaaattttattacaacagttattaatatacgatcaacagggattacccttgaccgaatacctctatttgtatgatcagtagttattacagcaattctactattactatctctc |
| 6be187338bb13b10478d04c3fc222c07af292b7b | Diptera | Drosophilidae | Stegana | Stegana psilolobosa | tctatcttcaggaattgctcatggaggagcatcagtagatttagcaattttttctcttcacttagctggaatttcttcaattttaggtgcagtaaattttattactactgtaattaatatacgtgcatcaggaatttctttagatcgaatacctttatttgtatgatctgtagtaattactgctttacttttattactatcttta |
| 1ac964fc37c57e2c643acf8fc8843b61058c836a | Diptera | Empididae | Hilara | Hilara hybrida | cctttcatctggaattgctcatggtggtgcttctgttgatttagcaattttctctcttcatttagcaggtatttcttctattttaggggctgtaaattttattactacagtaattaatatacgtacccctggaatttcatttgaccgaatacctttatttgtttgatcagtagtaattacagccatcttattattactttcttta |
| 1093c2b115fcecf8c8f4c6f31fc75116c7c8f139 | Diptera | Ephydridae | Ditrichophora | Ditrichophora occidentalis | tttatcatcagtaattgctcatggaggagcttcagttgatttagcaattttttctttacatttagctggaatttcttcaattttaggagctgtaaattttattactaccgtaattaatatacgatcaacaggaattacatttgatcgaataccattatttgtttgatcagtagtaattactgcattactacttttattatcatta |
| fd303f53e85c16842b9f9d9e759b2df8d6240420 | Diptera | Ephydridae | Parydra | Parydra paullula | gctatctgctggtattgctcatggaggagcttcagttgatttagctattttttcattacatttagcaggaatttcttcaattttaggagcagtaaattttattacaacagtaattaatatgcgagcaactggaattacttttgatcgaatacctttatttgtatgatcagtagttattactgctttattacttttattatcatta |
| 700bebdd3538598b388e262860258e0650c3013a | Diptera | Fanniidae | Fannia | Fannia intermedia | tctgtcttctaatattgctcacggaggagcttcagttgatttagctattttttctcttcatttagctggaatttcatctattttaggagctgtaaattttattactactgtaattaatatacgatctactggaattacatttgatcgaatacctttatttgtatgatcagtagtaattacagctttattattactgttatcatta |
| 87357c3b7927cc2d4b9aef962cda8233e982b9c0 | Diptera | Fanniidae | Fannia | Fannia ringdahlana | cctatcatctaacattgcacacgggggggcatcggtagacctagccattttctctttacatttagctggaatttcttcaattttaggagcagtaaattttatcaccactgtaattaatatacgagctacagggatttcatttgatcgtatacctttatttgtttgatctgtagtaattacagctttattattacttttatcattg |
| c29be6e79b5dd288e77ecafa6fe2b7f0e5552696 | Diptera | Fanniidae | Fannia | Fannia spathiophora | actatcatctaatattgctcatggaggtgcatccgtagaccttgctattttttctttacatttggctggaatttcatctattttaggagcagtaaatttcattacaactgtaattaatatacgagctacaggaattacatttgaccgaatacctttatttgtttgatctgtagtaattacagcactactattacttttatcatta |
| 63c11723c334ead9142a4bd1d24f31ca4a5fca54 | Diptera | Hybotidae | Drapetis | Drapetis completa | tctttcatcaggtatcgcccatagaggagcttctgtagatctagctattttttctttacatttagcagggatttcttctattttaggggccgtaaattttattacaacagtaatcaatatacgatccccaggaattacttttgatcgaatacctttattcgtatgatctgtagttattacagcacttcttcttttattatctcta |
| 38b24c89b522d81e81a1bf6a93d5c81fbc27099d | Diptera | Hybotidae | Tachydromia | Tachydromia luteicornis | cttatcgtcaggaattgctcacggaggggcatctgtagacttagcaattttttccctacacttagctggtatttcttcaattttaggagctgtaaattttattaccactgtaattaacatacgttctataggaattactcttgaccgcatacctctatttgtctgatcagtagtaattacagcaattttacttttattatctcta |
| 08bfe820b8a7969df9329187959aabe68bcf9886 | Diptera | Micropezidae | Micropeza | Micropeza thoracicum | tttatcatctggaatcgctcatggaggtgcatcagttgatttagctattttctcattacacttagcaggtatttcttctattttaggagctgtaaattttattacaactgtcattaatatacgatctacaggaattacttttgatcgaatacctttatttgtatgagcagtagtaattactgctttattactacttttatcttta |
| 055b07caaa59d212de2864990ed52a24c0b020ed | Diptera | Milichiidae | Neophyllomyza | Neophyllomyza acyglossa | cttatcttctattattgcacacggaagtgcttcagtagatttagcaattttttctttacatcttgcaggaatttcttcaattttaggagcagtaaattttattactaccgttattaatatacgatctacaggtattacgtttgaccgaatacctttatttgtttgatctgtagttattacagctttattattacttttatcacta |
| c252b71eb43d37773d102ecc38d22d691822b4d1 | Diptera | Muscidae | Gymnodia | Gymnodia humilis | tttatcatctaatattgctcatggaggagcttctgtagacttagctattttttcattacatttagcaggtatttcttctattttaggggctgttaattttattactactgtaattaatatacgatcaacaggaattacatttgatcgaatacctttatttgtttgatcagtagtaattacagctttattacttcttttatcttta |
| eaf9d51920b6ec49e4037703a96e4ab149808554 | Diptera | Mycetophilidae | Brachycampta | Brachycampta alternans | tctttcttcaacaattgctcatgctggtgcatctgttgatctagcaattttttctttacatttagcaggaatttcctcaattttaggagctgttaattttattacaactattattaatatacgagctccaggaatttcttttgatcgaatacctttatttgtttgatcagttcttattacagctgttctcttacttttatccctt |
| 3894db3354e7ea8b09f3a83ccc6795e88854f6a7 | Diptera | Mycetophilidae | Coelosia | Coelosia silvatica | attatcttctactattgcacatgctggagcctctgtagatttagctattttttctttacatttagcaggaatttcttcaattttaggagcagtaaattttattactactattattaatatacgagctcctggaatttcatttgatcgaatacctttattcgtatgatcagtattaattacagctattttattattactatctctt |
| 0ae28d365365cf15cf83aa49d272a1097f5b11f7 | Diptera | Mycetophilidae | Leia | Leia winthemi | tttggcatcaacaattgctcatgctggagcatctgttgacttagcaattttctctctacatcttgccggaatttcatctattttaggagcagtaaattttattaccacaattatcaatatacgagccccaggaatatcatttgatcgattacctttatttgtctgatctgtattaattactgcaattttacttttattatctctt |
| 7565685219b7eb8ad096588c98f6cbce99431a10 | Diptera | Mycetophilidae | Megophtalmidia | Megophtalmidia crassicornis | cctcgctgcaggaattgcccatgcaggctcctctgttgatttagctattttttctttacatttagctggtatttcttcaattttaggagcagttaactttattacgacaattattaatatacgagcccctggaatttcttttgaccgaatacctctatttgtttgatctgtcttaattactgctattttacttcttctttcttta |
| e8bdbc61d8d0516d661d4ecd2464d141be09e7cc | Diptera | Mycetophilidae | Opsion | Opsion nigra | tctttcttcaactattgcacatgcaggtgcttcagtagatttagctattttttctcttcatttagcaggaatttcctcaattttaggagctattaattttattacaacaattattaatatacgagctacaggaattacatttgatcgaatacctttatttgtttgatcagtatttattacagctattttattactattatcctta |
| 7b5adbb24d696914451423041af2c76ac0ad11e0 | Diptera | Phoridae | Aenigmatias | Aenigmatias lubbocki | cctttcatctaatattgctcataatggaccttccattgatttagtaattttctctcttcatttagctggaatctcctctattttaggatccattaattttatttctactattattaacatacatcccacaaaaataacactagaccgaatatctttatttatttgatctgtaataattacagctattttattacttttatcttta |
| 21d929fc0d2d419cd326a3e77f2ba18cab6bc6a6 | Diptera | Phoridae | Megaselia | Megaselia abdita | tttatcttctagaattgctcatagaggagcttctgtagatttagcaattttttcccttcatttagccggtatttcctcaattttaggagcagtaaattttattactacaatcattaatatacgatcagcaggtattacatttgatcgaatacctttatttgtatgatcagtaggaattactgctttactactattattatcactt |
| 0ae115ff3d23f296ea840d2208918b39b598701f | Diptera | Phoridae | Megaselia | Megaselia aculeata | attatcttctagaattgcacataatggagcatcagttgacttagcaattttttctttacatttagccggaatttcaagaattttaggtgctgtaaattttattacgaccattattaacatacgatcaacaggaattacttttgaccgaatacctttatttgtatgatctgtaggtattacagctttattattattgttatcatta |
| 29c2399f40d04c17915cdc5c5d3f827d0657ced5 | Diptera | Phoridae | Megaselia | Megaselia albicans | tttatcttctagaattgctcatagaggagcttcagttgatttagctattttttctcttcatttagcaggaatctcatcaattttaggagcagtaaattttattacaactattattaatatacgatcttcaggaattacttttgatcgaataccattatttgtttgatcagtaggaatcactgctcttcttttattattatcgcta |
| 85618381fa670d874c247bc114daff65f276c867 | Diptera | Phoridae | Megaselia | Megaselia bifida | actttcatctagaattgctcatagtggagcttctgttgatttagcaattttttctcttcatttagctggaatttcttcaattttaggtgcagtaaattttattactacaattattaatatacgatcttcaggtattacatttgaccgtatacctttatttgtttgatcagttggtattaccgctcttcttttattattatctctc |
| 86f76b3f5589892cbc12e5d3cacd0e71b7061e66 | Diptera | Phoridae | Megaselia | Megaselia drakei | actttcttctagtattgcccacagaggagcttccgtcgacttagctattttttcccttcatttagcaggtatttctagaattttaggagctgtaaattttattacaactattattaatatgcgatcttcaggaattacttttgatcgaatacctttatttgtttgatctgttggaattactgctctccttttattactttctctt |
| 648ba466edc58b933a5fd4f61b36ce453b11320b | Diptera | Phoridae | Megaselia | Megaselia haraldlundi | tctttcatcttcaattgcacatagaggagcttctgtagatttagctattttttctttacatttagcaggaatctcatcaattctaggagcagtaaactttattacaacaattattaatatacgttcttcaggtattacatttgatcgtataccattatttgtttgatcagttggtattacagctcttcttttattattatcttta |
| d4a61e488d7024971e9ad694da8cbe150b063bbd | Diptera | Phoridae | Megaselia | Megaselia hibernans | tttatcctctaacatcgcacatagaggggcatcagtagatttagcaattttttctcttcatttagcaggaatttcttcaattttaggagctgtaaattttattacaacaattattaatatgcgatcatcagggattacttttgatcgaatacctttatttgtatgatcagtaggaattactgctcttctacttttactttcactt |
| 3dee6ee2858042abe63273695cc12e2c278e94ae | Diptera | Phoridae | Megaselia | Megaselia ignobilis | tttatcctctagaattgcccatagaggggcttctgttgatttagctattttttcgcttcatttagcaggaatttcaagaattctaggagctgtaaattttattaccactattattaatatacgatcttctggtattacctttgaccgaatacctttatttgtttgatctgttggaattactgctcttcttttacttttatcatta |
| 447bb1d3d45c333219961fa473f5139ba2d7af4b | Diptera | Phoridae | Megaselia | Megaselia rufifrons | attatcttctaatattgcccatagaggatcctcagtagatttagcaattttttctttacatttagcgggtatttcttcaattctaggagcagttaattttattaccacaattattaatatacgatcttcaggaatttcttttgataaaataccattatttgtttgatcagttggaatcacagctttattactattgttatcttta |
| ecf1e0e20f8293131630335bd6ff16aa44e6bcda | Diptera | Phoridae | Megaselia | Megaselia sp. BOLD:ACX6055 | tctttcttctagtattgctcatagtggagcctcagttgatttagctattttctcccttcatctagctggtatttcttcaattttaggagctgtaaattttattacaacaatcattaatatacggtcatctggaattacatttgatcgaatacctttatttgtttgatcagtaggaatcactgctcttctattattattatcttta |
| ed62b90811f282fe72254e3ab2c6a16033efed0a | Diptera | Phoridae | Megaselia | Megaselia sp. BOLD:AAG3266 | actttcttctagaattgctcatagaggggcttcagttgatttagcaattttctctttacacttagctgggatttcttcaattttaggtgctgtaaattttattactacaattattaacatacgatcttcaggaattacttttgatcgaatacctttatttgtttgatcagtaggaattactgctcttttattacttttatcttta |
| b57d9e1d6ba531f4749784796f41728aacbd6925 | Diptera | Phoridae | Megaselia | Megaselia speiseri | cttatcttcaagaattgcacatagtggagcttctgttgatttagcaattttctcattacacttagcaggaatttcttcaattttaggtgcagtaaatttcattaccactattattaatatacgatcatcaggtattacatttgatcgtatacctttatttgtatgatcagtaggaattacagctttattacttttattgtcatta |
| 43033c06c0d72c9517dad09447bf804308331fe6 | Diptera | Phoridae | Megaselia | Megaselia subfuscipes | attatctgcaagaattgctcatagaggagcttctgttgatttagcaattttctctctacatctagcaggtatttcttctattttaggagcagtaaactttatcactacaattattaatatacgttcatcagggattacttttgatcgaatacctttatttgtttgatctgtaggaatcactgctcttttactattactatctcta |
| 0f7455cc394e095246b6c30f42734fb48868fbaf | Diptera | Phoridae | Megaselia | Megaselia verralli | tctttcttcaagaatcgctcatagaggagcctctgttgatttagcaattttttcacttcatttagctggaatctcaagaattcttggagcagtaaattttattactacaattattaatatacgatcttcaggtattacatttgaccgaatacctttatttgtttgatcagtaggtatcacagccttacttttattactctcctta |
| f03536c4fc12a3e731a6d5506bffd582dd869c23 | Diptera | Phoridae | Megaselia | Megaselia vestita | cctttcttctaatattgcacatagaggtgcatcagtagatttagcaattttttctcttcatttagcaggaatttcttcaattttaggagctgtaaattttattacaacaattattaatatacgatcaacaggtatttcttttgaccgtatacctttatttgtttgatcagtaggaattactgctcttcttttattattatctctt |
| fe6375ebc4d13d5898c4374ba0e8570edb9a2426 | Diptera | Phoridae | Phalacrotophora | Phalacrotophora delageae | tttatcctctagaattgctcatagaggagcttctgttgatctagcaattttctctctacatctagcaggaatttcctctattttaggagcagtaaattttattacaacaattattaatatacgatcatctggaattacatttgaccgaatacctttatttgtatgatcagtaggaattactgcccttcttcttcttctttccctt |
| d1f4a63e1e66995ccd1db0817aab4acad79f3195 | Diptera | Piophilidae | Piophila | Piophila nigriceps | tttatcttctgtaattgcacatggaggagcttctgtagatttagctattttctctttacatcttgctggaatttcttcaattttaggagctgtaaattttatcacgactgtaattaacatacgatctacaggaattacatttgaccgaatacctttatttgtttgatcagttgtaattactgctcttttattactattgtcactt |
| cb87397e15d94e62728efa271446b96bce38c213 | Diptera | Pipunculidae | Cephalops | Cephalops hardyi | tttatcttctaatattgctcatggtggagcctctgtagatttagctattttttcattacatctagctggaatttcatctattttaggagcagtaaattttattacaactgtaattaatatacggtcaaaaggaatatcttttgaccgaataccattatttgtgtgagctgtagctattacagctttattattattattatcttta |
| 6d6bf2c8b311586b26e9af39f28f91d7d9a653c6 | Diptera | Pipunculidae | Jassidophaga | Jassidophaga fasciata | attatcttcaacaattgctcatggaggtgcatcagttgatttagcaattttttcacttcatttagctggaatttcttcaattttaggagcagtaaattttatcactacagtaattaatatacgttcaacaggaattaattttgatcgaatacctctatttgtatgatcagtaattattacagcattacttttacttttatcatta |
| de5dc97aa0e90eac03547a016d0c3d5305777584 | Diptera | Platypezidae | Seri | Seri obscuripennis | cctatcttcaattattgctcacggtggggcttcggtagacttagctattttttctttacatttagctggaatctcttctattttaggagctgtaaactttattactactgttattaatatgcgagcagttggaatctctttcgaccgaatacctctgtttgtttgatcagttgttattacagctattcttttactactatcttta |
| a18e4efa42d0a919b2690cf30b2017b585a959a5 | Diptera | Psilidae | Psila | Psila sp. BOLD:AAF9710 | actatcatcaattattgcacatggaggagcttcagttgatttagcaattttttctttacatttagctggaatttcttcaattttaggagcagtaaattttattactacagtaattaatatacgatctacaggaattacatttgatcgaatacctttatttgtgtgatctgtagtaattactgctttattattattgttatcatta |
| 4f77fd248d9798008cc1adb1f46f12068fca12ba | Diptera | Psychodidae | Psychoda | Psychoda alternata | tctttctagattaatttctcatggcggaccttcagttgatttagctattttttctcttcatttagctgggatttcttccattttaggggcagtaaattttattactacaattattaatatacgatcaattgggattacatttgaacgaatacctttatttgtctgatcagtattaattactgcagttttattattactttcatta |
| 497d3dc6e0f2904ce618d2fca115332477ab7c56 | Diptera | Psychodidae | Psychoda | Psychoda lobata | tctttctagattagtctctcatggaggcccttcagttgatctggctattttttctttacacttagctgggatttcctcaattttaggggctgtaaattttattactacaattattaatatacgatctattggaattactttcgaacgaatacctttatttgtttgatctgttttaattacagcggtattacttcttctttcatta |
| cbd46ef20fbf522e406a59bdc1c72bbfc9a5b866 | Diptera | Psychodidae | Psychomora | Psychomora mycophila | tttatctagattaatttctcatggaggcccttcagtagatttagctattttctctttacatttagcggggatttcttcaattttaggagcagtaaattttattactacaattattaatatacgatcaatcgggattacatttgaacgaatacctttatttgtttgatcagttttaattactgctgttttattattattatcatta |
| 2e10b547ab4cc898ffa9c7d2746957b3dd78e4b8 | Diptera | Scatopsidae | Scatopse | Scatopse halterata | tttatcagcaggaattgctcatgggggagcttccgttgatcttgctattttttcattacatatagcaggtatttcttcaattttaggggctgtaaattttattacaactgtaattaatatacgatctccaggaattacttttgatcgaatacctttatttgtttgatcagttgttattactgctattttattacttttatccttg |
| 098d28e87d2c2a76f2674888332464b1d152a717 | Diptera | Sciaridae | Bradysia | Bradysia fenestralis | tttatcatcaacaatttctcattcgggtgcatcagttgatttatcaattttttcccttcacttagcaggtatttcatctattttaggggcagtaaattttatttcaactattattaatatacgggccccaggtatatcatttgataaaatacctttatttgtttgatcagtattaattacagctatccttttattactatcatta |
| 7fc1d581af71220ec5fefc70a00c4ce6c54fd7e0 | Diptera | Sciaridae | Bradysia | Bradysia giraudii | actttcttcttcaattgctcatagaggagcctctgttgatttagccatcttttctctacatctagctggaatctcatctattttaggagccgtaaattttatcactactattattaacatacgatcatcagggattacttttgaccgtatgcccttatttgtttgatcagtaggaattactgctattttacttttattatcactt |
| 11557a289006cd3afea21c6dd70b5bfaa05b63b0 | Diptera | Sciaridae | Bradysia | Bradysia longistylia | tttatcctcaacattatctcattctggggcttcagttgacttatcaattttttctcttcatttagctggtatttcttccattctaggggctgtaaattttatttcaaccattattaatatacgaacacccgggataagttttgataaaatacctttatttgtctgatctgtattaattacagctatcttattacttctatcatta |
| 111c5983097d62cc573c3136e73e28a5b12c9015 | Diptera | Sciaridae | Bradysia | Bradysia pallipes | tctttcttctaccttgtctcactcaggagcgtcagtagacctttcaattttttcattacacttagcaggtatttcatcaattctaggagcagtaaattttatttctactattattaatatacgggctcccggaatatcatttgataaaataccattatttgtatgatctgttttaattacagctgtattattattattatcccta |
| 45f5519c08f6b64115933f6622bb38439776b68e | Diptera | Sciaridae | Bradysia | Bradysia peraffinis | tctatcttctacaatctcccactctggggcatcagttgatctttcaattttctctcttcatttagctggaatttcttcaattttaggggctgtgaattttatttcaactattattaatatacgaacacccggtatatcatttgataaaatacccctttttgtatgatccgtcttaattacagctattttattattattatctctt |
| e0caae20dafb1234809978e5f9012fe2bcb39f9e | Diptera | Sciaridae | Bradysia | Bradysia polonica | tttatcttcaactttagcacattctggagcttcagttgatttatcaattttctctcttcatctagctggaatttcctctattttaggagcagtaaactttatttccacaattatcaatatacgagcaccagggataacttttgataaaatatcattatttatttgatcagttcttatcaccgcaattcttcttcttctttctctt |
| cb8a9187ed245dbde4634210fccfa6f63b8519bc | Diptera | Sciaridae | Bradysia | Bradysia procera | tttatcttcaacaatttctcattcaggagcatcagtagatttatcaattttttcattacacttagcaggaatttcttctattttaggggcagtaaattttatttctacaattattaatataaaggcacccggtttatcatttgataaaattcctttatttgtttgatctgttttaattacagctattttacttttattatcatta |
| a52ae7ecefb59ee457a9d02ce612577a3273df28 | Diptera | Sciaridae | Bradysia | Bradysia scabricornis | tctttcttctaatatttctcattcaggcgcatcagttgatttatctattttttctttacatttagctggtatttcatcaattttaggagcagtaaattttatttccacaattattaatatacgcacaccaggaatatcttttgataaaatgcctttatttgtatgatctgtattaattaccgcagttctattactattatctcta |
| 78294f72d7edfce44986832722c7c769d9125c6b | Diptera | Sciaridae | Bradysia | Bradysia strenua | tctatcatcaacactctcacattcaggggcatctgtagatttatctattttttctctgcacttagcaggtatttcttcaatcttgggagctgtaaacttcatttcaactattattaatatacgagccccaggtatatcattcgataaaatacctttatttgtttgatcagtattgattacagctgtattattgttattatcatta |
| 8f1d812052b9543854c0a196aa31d2f862ba91e6 | Diptera | Sciaridae | Bradysia | Bradysia urticae | actttcttcgacaatttctcattcaggagcatcagttgatctatcaattttttctttacatttagcaggtatctcttctattttaggtgcagttaacttcatttctactattattaatatacgagcccctggaatgtcctttgataaaatacctttatttgtatgatccgttttaattacagcaattcttttattattatcttta |
| 13fc2d2f15da4efec12efbc839fb5f8e94b5b41d | Diptera | Sciaridae | Bradysia | Bradysia vagans | cctttcatcaacactatcacactcaggggcttcagttgacttatcaattttttccctccatctagcaggtatttcatcaattttaggggctgtaaattttatctcgactatcattaatatacgggcccctggtatatcttttgataaaatacctttatttgtttgatctgttttaattaccgccgtattactattattatcttta |
| 5340c19623045b012cd5450347df0742b2f46367 | Diptera | Sciaridae | Camptochaeta | Camptochaeta cursor | attatcatcaacactatctcattctggtgcttcagtagatttatctattttttctctacatttggctggtatttcttcaattttaggagctgtaaattttatctctactatcattaatatacgaacccctggtatatcatttgataaaataccattatttgtttgatcagtattaattacagccgtattgttgttattatcttta |
| b9f233a66a52c66606728c5ce5331c2c3061d44e | Diptera | Sciaridae | Camptochaeta | Camptochaeta furcata | attatcttcgacattatcccattcaggagcatctgtagatttgtctattttttccctacatttagctggaatttcttccattttgggggcagtaaattttatttccacaattattaatatgcgggccccaggcatattgtttgacaaaatacctttatttgtatgatcagtattaattactgcaattctattattgttatcttta |
| d38653798e74c2649e6b9c5b72b0cf361f7dfb07 | Diptera | Sciaridae | Camptochaeta | Camptochaeta vivax | attatcatctactatatcccattcaggagcttctgtagatttatctattttttcattacacttggcaggaatttcctctattttaggtgcagtaaattttatttctacaatcattaatatacgagcccccggtatatcttttgataaaatacccctctttatttgatcagtattaattacagctattttattattattatcatta |
| cb8c29f49a5bdc3009cc4a216ef0866857124485 | Diptera | Sciaridae | Claustropyga | Claustropyga aperta | ttatcttcaactctttcccattcaggagcttcagtagacttatcaattttttctttacatctagctggaatttcatcaattttaggagcagttaattttatttctactattattaatatacgtgctccagggatatcttttgataaaatacctttatttatttgatcagtattaattacagccatcttactacttttatcttta |
| ea5e8704166bd0965d47e2d64a90f8dc52d8e7c8 | Diptera | Sciaridae | Corynoptera | Corynoptera anae | attatcttcaactttatctcactcgggggcctcagttgatttatcaattttttctcttcatttagcaggtatttcatcaatcctgggggcagtaaattttatttctacaattattaatatacgggccccaggtatatcatttgacaaaatacctttatttgtatgatcagtattaattacagctgtacttttacttttatcatta |
| d71ccf8ebf4010edd75902b02c44bfc735b9ee1d | Diptera | Sciaridae | Corynoptera | Corynoptera bicuspidata | cttatcatcaactctatctcattcaggagcatcagttgatctatcaattttttcattacatttagcaggaatttcttctattcttggcgctgtaaattttatttctactattattaatatacgagcccctggtatatcttttgataaaataccattattcgtttgatcagttttaattacagctattttactacttttatcatta |
| 5ac7eb592c52f0cbb0de8bd8fd778bb33732686b | Diptera | Sciaridae | Corynoptera | Corynoptera cuniculata | attatcttcaaatttatcccattcaggagcctcagttgatttttccattttttctcttcatttagcaggtatttcatcaattttaggggcagtaaattttatttctacaattattaatatacgagctcctgggatgtcatttgataaaatatccctatttgtatgatcagttttaattacagctattttattactattatctctg |
| 9e8e51e8e8e35a4c16b2aa733279ddb53c241e86 | Diptera | Sciaridae | Corynoptera | Corynoptera dentata | tctttcttctacattgtctcattctggtgcctctgttgatttatcaattttttctttacatttagctggtatttcctctattttaggggctgtaaattttatttctactattattaatatacgggcccccgggatgttttttgataaagtttcattatttgtttgatctgttcttattactgctgttttattattactttcttta |
| c248572dfb8296d93413380459eb0a33be4fbae3 | Diptera | Sciaridae | Corynoptera | Corynoptera flavicauda | ttatcatcaacattatctcattcaggggcatctgtagatttatctattttttctcttcatttagccgggatttcgtcaattttgggagctgtaaatttcatctcaacaattattaatatacgggcacccggaatattatttgataagatacctctatttgtatgatcagtactaattacagcaattttattattactatctctc |
| d8783c83214e6f36dc3644451a041af513624400 | Diptera | Sciaridae | Corynoptera | Corynoptera inundata | tttatcttctactttatctcattctggtgcatctgttgatttatcaattttttctcttcatttagcaggaatttcttcaattttaggagctgtaaattttatttcaactattattaatatacgagctcctggtatatcatttgataaaatacctttatttgtatgatcagttttaattactgctattttattacttttatcatta |
| 246ac0c05314a897075e1ace1acda22feeafae1e | Diptera | Sciaridae | Corynoptera | Corynoptera minima | cctttcctctactttatctcattcaggagcatcagtcgatttatccattttttcccttcatttagctggtatttcttcgattttaggggcagtaaattttatttcaaccattattaatatacgaacgccgggaatatccttcgataagacacccttatttgtgtgatcagtattaattactgcggtattattactactatcttta |
| 7d5a444288f2aa38b73ce4dd069e5be439430c62 | Diptera | Sciaridae | Corynoptera | Corynoptera subcavipes | ctatcatctactatttcccattccggagcttctgttgacttatcaattttttccctccatttagcaggtatttcatctattctaggggctgtaaactttatttcaacagttattaatatacgagcccccggcataatatttgataaaatacctctatttgtatgatcagtattgatcacagcaattttgttattgttatcccta |
| cbeef04d758d28933881c242d72bbd2aee136ffa | Diptera | Sciaridae | Corynoptera | Corynoptera trepida | tttatcttcaactttatcccattcaggagcctctgtagatttatctattttctctcttcatttggctggtatttcttctattttaggggctgtaaattttatttctactattattaatatacgcgccccaggaatattatttgataaaatacctttatttgtttggtccgttctaattacagccattttattattattatccttg |
| 24158a80dd4bc4a1992aa0e517c17a60d3b95816 | Diptera | Sciaridae | Corynoptera | Corynoptera umbrata | tctttcatctaatttatctcatgcaggagcatcagttgatatatccattttttccctacatttagcaggaatttcctctattttaggggccgtaaattttatttctacaattattaacatgcgcaccccaggaataatgtttgatataatacccttatttgtgtgatcggtattaatcactgctattcttctacttttgtccctc |
| f3b988d2a399133e683d0b4be1affdaa472c1dda | Diptera | Sciaridae | Cratyna | Cratyna crassistylata | tctatcatctactctttcccactcaggagcttcagtggacctatctattttctctttacatttagcaggaatttcttcaattttaggggctgtaaattttatttcaaccattattaatatacgagccccaggcatatcttttgataaaatacctttatttgtgtgatcagtgttaattacagccattttattattattatctctt |
| 21e492919d204b35dc2d99cf2cffc7ef7da9ab18 | Diptera | Sciaridae | Cratyna | Cratyna uliginosoides | tttatcctctactttatctcattcaggagcctctgtagatctctctattttttcactccatttagcaggaatttcttctattttaggggcagtaaactttatttctactattattaatataaaaactcctggaataacttttgataaaatacctttatttgtctgatctgtttttattacagctatcttattactcttatcttta |
| 8e7e05bc903d31f2a432b47c55aff9db1e47d028 | Diptera | Sciaridae | Dichopygina | Dichopygina nigrohalteralis | cctatcttctactttatcccactcaggggcatctgtagacctatcaattttttctttgcacttagcggggatctcttcaattttaggcgctgtaaactttatttcaactattattaatatacgggcccctggcatatcttttgataaaatccccctatttgtttggtctgtattaattactgctattttactattattatccctc |
| 3b0de06866283a3ce89c3bf48a7d694f512513b1 | Diptera | Sciaridae | Epidapus | Epidapus alnicola | tctttcatctactttatctcattctggttcttccgtagatttatctattttttctttacatcttgcgggtatctcatctattttaggggcagttaattttatttctactattgttaatatgcgggcgtttgggatgttatttgataagatttctttatttatttgatctgtattaattactgctattttattgttgctttcttta |
| e9ad1411a24ca6f82d2620537effd53b9362ce40 | Diptera | Sciaridae | Epidapus | Epidapus ignotus | tctatcagctaatttatcccattcaggagcatcagtagatttatcaattttttctttacaccttgctggaatttcatccatcctaggggcagtaaattttatttctactattattaatatacgaactagaggtatacaatatgataaaactcccttatttgtttgatcagtattaattactgcaatcttactattattatcttta |
| 0588cac6b718188e4ff14d2b66a3275b1b6f7adf | Diptera | Sciaridae | Lycoriella | Lycoriella acutostylia | cctatcttctactctatctcactcaggagcttctgtagacttatcaatcttttctttacatttagcgggaatttcctcaattctaggagccgtaaattttatctcaacaattattaatatacgagccccaggaatatcctttgacaaaattcctttattaatctggtcaatcctaattactgcaattttattattgctatccctt |
| 4651f8e442d16d10ff15417295234f8bcc882d82 | Diptera | Sciaridae | Lycoriella | Lycoriella conspicua | tttatcttcaactttagcccattcaggagcttcagtagatctatcaattttttcattacatttggcaggaatttcatcaattttaggggcagtaaattttatttctacaattattaatataaaagcccctggaataacttttgacaaaatgcctctttttatttgatcagttttaattactgctatcttattattattatctcta |
| 81476a27251250a065eafbd2d4cbc469f40cb0b7 | Diptera | Sciaridae | Lycoriella | Lycoriella inflata | tttatcctcttctttatctcattctggggcttctgtagacctatcaattttttctcttcatttagccggtatctcatcaattctaggtgcagtaaatttcatttctacaattattaacatacaagccccaggaatatcctttgataaaataccattattcatttgatctgttttaattactgctattttattattattatctctc |
| 899f93d9eb589281ab329144ee9016f11f06d8bb | Diptera | Sciaridae | Lycoriella | Lycoriella ingenua | tttatcttctactgtatcccactcaggggcttctgtagacttatctattttttctttacatttagccggaatttcatcaattttaggtgctgtaaatttcatttctactattattaatatacgagcccctggaataacctttgataaaatacccttatttatttgatctgtaggaattactgctactttacttctattatctcta |
| b536d6b938fda12ebc82a3cada6d827b73f6d96e | Diptera | Sciaridae | Lycoriella | Lycoriella sativae | cctatcttctactttagctcattcaggggcttccgtagatttatctattttttctttacatttagcgggtatttcctcaatcttgggggcagtaaattttatttccactattattaatatacgagcccctggaatatcttttgataaaataccattatttatttgatcagttttaattactgcaattcttcttcttctatcccta |
| 7a68c6e3ee4620534c357c88ad94dd04178cfd4e | Diptera | Sciaridae | Lycoriella | Lycoriella weberi | tctttcttcaactttatctcattcaggagcttccgtagatctttctatcttctctttacatttagctggtatctcttcaattttaggggcagtaaactttatttctactattatcaatatacgtgcaccaggcatatcatttgataaaatacctttatttatttgatcagttttaattacagctattcttctattgttatcttta |
| b92fbdd4a94ed92ae942e6963ff35f21b80b79a0 | Diptera | Sciaridae | Scatopsciara | Scatopsciara brevicornis | cctatcttcaactctatcccactcaggggcctcagttgacttatcaattttttcccttcacttagcagggatctcttctattctaggggcagtaaattttatttctaccattattaatatacgggcccctggcatatcttttgataagatacctttatttgtctggtccgttcttattactgcagtccttttactactctctctc |
| 36c3ee26b4fed439175425eb93194aeddbf320cf | Diptera | Sciaridae | Scatopsciara | Scatopsciara calamophila | tctttcgtctactttatcccactcgggggcatcagttgatctatctattttttctcttcatttagctggaatctcttcaattttaggggcagtaaactttatttctaccattattaatatgcgagccccagggatatctttcgataaaatacccttatttgtatgatcagtacttattactgcagtacttctattattatccctt |
| 35f5d32ebbf93bfb7975de58507a0b5961f6249a | Diptera | Sciaridae | Scatopsciara | Scatopsciara dentifera | tctttcatccacgttagctcattcaggggcttctgttgacctatctattttctcccttcatttagcaggaatttcttcaattcttggggctgtaaactttatttccactattattaatatacgagcttctgggatatcatttgacaaaatacccctatttatttgatccgtttttattacagccattttattgcttctttcatta |
| 6cdaeed99932d1f3bfbc331b3b989b7c4f72fe02 | Diptera | Sciaridae | Scatopsciara | Scatopsciara edwardsi | attatcttcttctattgctcactcaggggcttctgttgatttatctattttttccctacacttagctggtatctcttctattttaggggcagttaattttatttcaactattattaatatacgagctcctggtatatcctttgacaaaatacctctatttgtatgatcagtattaattacagcaattcttttacttttatcttta |
| e2675ba8ba6d47098685f80e15f2615847ce6d62 | Diptera | Sciaridae | Scatopsciara | Scatopsciara multispina | tctttcttcaacactttcccattctggggcttcagttgatctttccattttttcccttcatttagcaggaatttcttcaattctaggagctgtaaattttatttcaactattattaatatacgagctcctggtatatcctttgataaaatacctttattcgtttgatctgtattaattacagcaatcctattattattatcatta |
| f03ac00566b4be8c68745355a6986f1abdaa1f27 | Diptera | Sciaridae | Scatopsciara | Scatopsciara pusilla | actatcttctactattgctcattcgggggcctctgttgatttatctattttttctttacatttagccggtatctcttctattttaggggccgttaattttatttctactattattaatatacgagccccaggtatatcatttgataaaatacctctattcgtttgatctgtactaattacagcaatcctattgcttttatctctg |
| 518d01284b9f33c10cbd5d6407c1c0f3043c6890 | Diptera | Sciaridae | Scatopsciara | Scatopsciara sibirica | tctatcttcaactttatctcattcaggggcttcagttgacctatctattttttcactccatttagcagggatctcttctattttaggggcagtaaattttatttccactatcattaatatacgggctccaggtatatcttttgataaaatgcctttatttatctgatcagtccttattacagcagtccttctattattatcccta |
| cdf942b42154606b53e959b639de5dace6870a60 | Diptera | Sciaridae | Scatopsciara | Scatopsciara vitripennis | actatcctctacattatcacattccggggcttcagtagatttatcaattttttctcttcatttagcagggatttcttcgatcctcggagccgtaaattttatttctactattattaacatgcgggcccccggaataacctttgacaaaatacctttatttgtctgatctgtattaatcacagcggttcttcttcttttatctctt |
| faaf197e05574881b1c243bccf923009258651e8 | Diptera | Sciaridae | Schwenckfeldina | Schwenckfeldina pectinea | tttatcatctactatttcacattcgggagcctcagttgatttatctattttttcccttcatttagccggtatttcctcaattttaggagctgtaaattttatttcaactattattaatatacgaattacaggaataacctttgacaaaatatctttatttgtctgatctgtatttattacagctattcttcttcttttatcttta |
| c8d9b0cdb4285eab5b6902744ee56ca363105240 | Diptera | Sciaridae | Sciara | Sciara atomaria | tctatcatcaactttatctcattcgggggcatctgtagatctctcaattttttctttacatttggcgggaatttcttctattttaggggctgtaaattttatttcaactattattaacatacgggcccctggaatatcttttgataaaatacctttattcgtatgatctgttcttattacagcagttttacttttattatccctc |
| 489f2a60ea0ed464bcea87de5e6c64671c6e26b9 | Diptera | Sciaridae | Sciara | Sciara concinna | tttatcatcaactctttctcattcaggggcatctgttgatttatcaattttttccctccatttagctggtatttcatcaattttaggggcagttaattttatttcaacaattattaatatacgagccccaggaataatatttgataaaatacctctttttgtatgatcagtattaattactgcaattctactacttctatcatta |
| 8cdf7eb0b8bd0522594007353e888457190d1d16 | Diptera | Sciaridae | Trichosia | Trichosia basdeni | attatcatcaactatttcacattctggggcatctgtcgatttatctattttttctcttcatcttgcaggtatttcttcaattttaggggcagtaaattttatttctacaattatcaatatacgagctccaggaatattttttgataaaatacctttatttgtttgatctgttttaattacagctattttattattattatcacta |
| 0e472ddcf228e2fe9288ae2247385f5ba6b78c0f | Diptera | Sciaridae | Xylosciara | Xylosciara heptacantha | tttatcatcaactctttcccattcaggagcttcggtagatttatcaattttttctcttcatttagcaggaatttcatctattttaggggcagtcaattttatctcaacaattattaatatgcgaaccccaggaataacctttgataaaataccgctttttgtatgatctgtctttattactgcaatcttattacttttatcttta |
| bcbecce6784f0098bd0c55ed30488ad19a05cfd9 | Diptera | Sciaridae | Xylosciara | Xylosciara misella | attatcatccacattatcccattcaggtgcgtcagttgatttatcaattttttcattacatttagctggtatttcatctattctaggagctgtcaactttatttcaacaattattaatatacgaacccctggtatatcttttgataaaatacctctttttgtttgatctgtattaattacagctattttattattattatcatta |
| 9f7d0af693429ea9bf92f0eae1b5de0919fe2ad6 | Diptera | Sepsidae | Sepsis | Sepsis neocynipsea | tttatcttctgggattgcccatggaggtgcatcagtagacttagctattttttcattacacttagcaggaatttcttctattttaggagctgtaaattttatcacaactgttattaatatacgatcaacaggaattacctttgaccgaatacctttatttgtttgagctgttgtaattactgccttattattacttttatcccta |
| 44f822b371fb482e80d27d66cf1893be7d62d895 | Diptera | Sepsidae | Zuskamira | Zuskamira inexpectata | tctttcatctgggatcgctcatggaggagcttcagttgatttagcaattttctctcttcatttagccggaatttcatcaattttaggggcagtaaattttattactactgtaattaatatacgatctaccggaattactttcgatcgaatgcctttatttgtttgagctgttgtaattactgctttattattacttctttcacta |
| 1492cb593bbf9cc86a10317d8c3f198d75e1a1e4 | Diptera | Sphaeroceridae | Ischiolepta | Ischiolepta intermedia | tttatcagcaggaattgctcatggaggagcatctgttgatttagctattttttcattacatttagctggaatttcttcaattttaggagctgtaaattttatcacgactgttattaatatacgatcaactggaattacatttgaccgaatacctttatttgtttgatctgttgtaattactgctttattattacttttatcatta |
| 6ca078d40b2c8df9425649c3c332ceb132e89668 | Diptera | Sphaeroceridae | Limosina | Limosina manicata | tctttcgtctaatattgctcatggtggagcttcagttgatttagcaattttttctttacatttagctggaatttcttctattttaggggctgtaaattttattactactgtaattaatatacgatcagtaggtattacttttgaccgaatacctttatttgtttgatcagtagtaattacagctcttttattattattatcttta |
| cf6d8c424e287b9f58ca875c0b72ebfec6595d84 | Diptera | Sphaeroceridae | Minilimosina | Minilimosina gemella | gctatcttctggtattgcacatggaggagcatcagtagatttagctattttttctcttcatttagctggtatttcttctattttaggggcagtaaattttattacaacagtaattaatatacgatctacaggaattacttttgatcgaatacctttatttgtatgatcagtagtaattactgctttattattattattatcttta |
| 6edd782c3ff98f32b99c0aff299dc745d300768b | Diptera | Sphaeroceridae | Minilimosina | Minilimosina parva | cctatcttctggtattgctcatggaggagcttctgtagatttagcaattttttcccttcatttagctggaatttcctctattttaggagcagtaaattttattactacagtaattaatatacgatctgtaggtattacatttgatcgaatacctttatttgtttgatctgtagtaattacagctttattactacttttatccttg |
| 4b3dfcf9d60fa3e5adbe905b143701790c72adde | Diptera | Sphaeroceridae | Opacifrons | Opacifrons humida | tttatcttctggaattgctcatggaggagcttcagtagatttagctattttttcattacatttagctggaatttcttcaattttaggagccgtaaattttattacaacagtaattaatatacgatctacaggaattacttttgatcgaatacctttatttgtatgatcagtagtaattactgctttattattacttttatcttta |
| 6eb4d5be0bddde0102af8542e91d4c7dcbe96e37 | Diptera | Sphaeroceridae | Pullimosina | Pullimosina pullula | tctttcttcagggattgcacatggaggagcatcagtagatttagctattttttctttacatttagctggaatttcttctattttaggagctgtaaattttattactacagtaattaatatacgatcaacaggaattacttttgaccgaataccattatttgtatgatctgtagtaattactgccttattattattattatcactt |
| f072f6250d58ac0867c489a89365dd93eead0692 | Diptera | Sphaeroceridae | Trachyopella | Trachyopella lineafrons | tctatcttctaacattgctcatggaggagcttctgttgatttagctattttctctcttcatttagctggaatttcctcaattttaggggctgtaaattttattacaactgtaattaatatacgatctacaggaattacattcgatcgaatgcctttatttgtttgatctgtagtaattactgcgttattacttttattgtcattg |
| d7cef18f5f2b8f368cb52f92d9b2c4ce931f2553 | Diptera | Syrphidae | Parasyrphus | Parasyrphus vockerothi | tctttctgctagaattgctcatggaggagcttctgttgatttagctattttttctcttcatttagctggtatatcttcaattttaggagcagtaaattttattactacagtaattaatatacgatctaatggactttcttatgatcgaatacctttatttgtatgatcagtagtaattacagctttattattacttttatcatta |
| 099625cdd7895b58cd9102ae62b8c77b2177e406 | Diptera | Syrphidae | Sericomyia | Sericomyia militaris | actttcatctagaattgctcatggaggagcttcagtagatttagctattttttctttacatttagctggtatatcttctattttaggtgcagtaaattttattacaacagttattaatatacgatcttctggaatttcttatgatcgaatacctttatttgtatgatcagttgttattactgctttattacttcttttatcatta |
| 5650179d8943832024ebe36e6bd92ad53c74bed2 | Diptera | Tachinidae | Gastrolepta | Gastrolepta anthracina | tttatcatctattattgcacatggaggagcttctgttgatttaacaattttttcattacatttagcaggtatttcatcaattttaggagcagtaaattttattacaacagtaattaatatgcgatctacaggtatttcatttgatcgaatacctttatttgtttgatctgttgttattacagctttattattacttttatcttta |
| 0a67203498c9d4df05a5de23c85fe356af7c99c0 | Diptera | Tachinidae | Siphona | Siphona nigricans | tttatcatctaatattgctcatagaggaacctcagtagatttagcaattttttctttacatttagcaggaatttcttcaattttaggagccgtaaattttattactacagtaattaatatacgatcaccaggtattacttttgaccgaatacctttatttgtttgatctgtagtaattacagctttacttttattattatcatta |
| 78a26da2886416c88876b3f45253a6f48506ad25 | Diptera | Tachinidae | Siphona | Siphona paludosa | tttatcatctaatattgctcatagaggaacttcagtagatttagcaattttttctttacatttagcaggaatttcctcaattttaggagctgtaaactttattactacagtaattaatatacgatcaccaggtattacttttgaccgtatacctttatttgtttgatctgtagcaattacagctttacttttattattatcatta |
| 55a7d2fd56101d1c8378025c415e36e779d26b24 | Diptera | Tephritidae | Campiglossa | Campiglossa farinata | cctttcatctatttcaactcatacaggagcttctgtagatttagctattttttctttacatttagcaggtatttcttcaattttaggagcagtaaattttattacaactgttattaatatacgatcaacaggaattacatttgatcgaatacctttatttgtatgagctgtagttttaactgctctcttacttttattatctcta |
| 67bc354f04f703abb9d7f9a58b35e0c15bcbe0d8 | Hemiptera | Aphididae | Allocotaphis | Allocotaphis quaestionis | attatcaaataatattgcacacaataatatttcagttgatttaactattttctctctacatttagcaggaatttcatcaattctaggagcaattaattttatttgtacaattttaaatataataccaaataatataaaattaaatcaaattcctctcttcccatgatcaattttaattacagcaattttattaattttatcttta |
| 9f9a4fc41f82e850fd3fd8fec10686dca6b6fb50 | Hemiptera | Aphididae | Aphis | Aphis neospiraeae | tttatcaaataatattgctcataataatatttcagttgatttaactattttttctcttcatttagcaggtatttcatcaattttaggagcaattaattttatttgtactattttaaatataataccaaataacataaaattaaatcaaattcctctatttccatgatcaatcttaattacagctatactattaattttatcttta |
| 394ff2fe8fb7e79d8af70b7182249df01e01ee31 | Hemiptera | Aphididae | Aphis | Aphis spiraecola | tttatcaaataatattgctcataataatatttcagttgatttaaccatcttctctcttcacctagcaggtatttcatcaattttaggagcaattaattttatttgtacaattcttaatataataccaaacaatataaaattaaatcaaatcccactatttccatgatcaatcttaattacagctatattattaattttatctcta |
| ae16a2c417511f29b1a0f57a3bd29eef0abb3290 | Hemiptera | Aphididae | Cinara | Cinara brauni | actatcaaataatattgcacataataatatttcagtagacttaactatcttttctctacatttagcaggaatctcatcaattttaggggcaattaattttatctgtacaattctaaatataatacctaataatttaaaacttagtcaaattccattattcccatgatcaattttaattacagccatattattaattttatcatta |
| 1678c65340d044da597f68fad3ad727cb64fac12 | Hemiptera | Aphididae | Hyadaphis | Hyadaphis passerinii | attatcaaataatattgcacataataatatttcagtagatttaactattttttctttacacttagcaggaatttcatcaattttaggagcaattaattttatttgcacaattcttaatataatacctaataatataaaaattaatcaaattcctcttttcccatgatcaatcctaattacagctattttattaattttatctcta |
| e2a050fe227f2b152da212d13ab891e053fa4410 | Hemiptera | Aphididae | Linosiphon | Linosiphon galiophagum | tttatcaaataatattgcacacaataatatttcagttgatttaactatcttttctttacatttagctggaatttcatcaattttaggagcaattaattttatttgcacaattctcaatataataccaaataatataaaattaaatcaaattcccctttttccttgatcaatcctaattacagctattttattaattttatcttta |
| 045c2c5e2597539927e86fd3dbc9d6e72bac8353 | Hemiptera | Aphididae | Lipaphis | Lipaphis pseudobrassicae | tttatcaaataatattgctcacaataatatttcagttgatttaactattttttctctacacttagcaggaatctcttcaattttaggagcaattaattttatctgcacaatcctaaatataatacctaataatataaaattaaatcaaattcctcttttcccatgatcaattttaattacagctattttattaattttatctcta |
| 02316f22e08c4faec1578ca3c959e02c7009d70a | Hemiptera | Aphididae | Schizaphis | Schizaphis graminum | attatctaataatattgctcataacaatatttcagttgatttaacaattttttccttacatttagcaggaatctcctcaattctaggagcaattaactttatttgtacaattttaaatataatacctaataatataaaattaaatcaaatcccattattcccttgatcaattttaatcacagctattttattaattttatcacta |
| 0b773750c895c330693e9f74e3f1f9aff27f4a84 | Hemiptera | Aphididae | Tuberculatus | Tuberculatus remaudierei | actatccaataatattgctcataataatatctcagtagatttaactatcttttccttacacttagcaggaatctcatcaattttaggagcaattaattttatttgtacaattcttaatataatacctaataacataaaattaaatcaaatccccctattcccttgatcaattttaatcacagcaattttattaattttatcctta |
| 43d4fe37c53cd603643f9a5662f851c1f4ecd42e | Hemiptera | Cicadellidae | Euscelis | Euscelis sordida | cctttcaagtaacattgcccattcaggagcaagagtagatttatctatcttctctctacacttagcaggaatttcttcaattttaggtgcagtaaatttcatttcaactgtaataaatatacgaacatcaggaatactgatagaccgcactcctctttttgtgtgatcagttcttattactgcaattcttttacttctgtcatta |
| 61b8dde6c79a46c29069b39f5b540259340566d8 | Hemiptera | Cicadellidae | Fagocyba | Fagocyba douglasi | cctttctagaaatatagcacactcaggatcaagtgttgatttaactattttttcgctacatttagcaggtatttcatcaattctaggagcagtaaattttatcacaacagtaataaacatacgacctatgaatataaaaatagaccgaacaccgctatttgtatgatcagtactcattacagcaatcctacttcttctgtcccta |
| d4d01fe32b6beeb842e4ed9a342fdb974cbb2447 | Hemiptera | Cicadellidae | Macrosteles | Macrosteles alpina | cctatctagaaatattgcacacgcagggccgagagttgatatatctatcttttccttacatttagctggtatttcttctattttaggggcagtaaattttattactaccgtaataaatatgcgccccacaggaatatctatagaacggacacctttatttgtatgatcagttttaattacagcagtcttgttacttttatcactc |
| bedd25d69d4bdb2bbf23d3e081e038cf2b873f1c | Hemiptera | Cicadellidae | Sorhoanus | Sorhoanus pascuellus | cctatctagaaatattgcacatgcaggagcaagagttgatatgtcaatcttttcccttcatttagctgggatctcctctatccttggggcagttaacttcatttcaacagtaataaatatgcgacctacaggaataacactagaccgaaccccactatttgtatgatctgtgttaatcactgccgtcctacttctattgtcttta |
| 433aa762ca7f4e2b055a2aad2d98ff73a290e19f | Hemiptera | Cicadellidae | Zyginidia | Zyginidia scutellaris | tctatcttctaatattgcccattcaggagcaagagtagatttagctattttttctcttcatctagccggaatttcttcaattctaggagcagtaaattttattacgaccgtaattaatatacgctcaattagattaacattagaccgaattcctttatttgtttgatctgttgtgattacagcaattttactactcctttcccta |
| bac5912832bc40317d7b0014d2cb9b3c5e9d6c9a | Hemiptera | Delphacidae | Nothodelphax | Nothodelphax consimilis | tctgtcaagaatcacttcacattcagggccctcagttgatctaacaatcttttcccttcacatcgctggtgttagttcaattataggagcaattaatttcatctcaacaattatcaatatgcattctaaaactatttcaatagaaaatttacccttattctgttgatcagttttaattacagctattctactactcctttcccta |
| d33fdb3f9a8e4ec299eeba6e6b908762c17e4cb0 | Hemiptera | Miridae | Monalocoris | Monalocoris amamianus | tttatcacataatatttcccataatggggcatcagtagatctagcaattttttccttacacttagcaggtatttcatcaatcttaggagcagtaaattttatttcaacaatcatcaatatacgagcaataggaataacagctgatcgtttaccattatttgtgtgatccgtaggaatcacggcattattattattattatcatta |
| 0269283a9458e3f6185a7d6878a4996e886b0c16 | Hymenoptera | Braconidae | Apanteles | Apanteles lacteicolor | attatcattaattttaggacatggaggtatatctgttgatttgggaattttttctttacatttagctggtgcttcatcaattataggggctgttaattttattaccaccattttaattatacgaacaaatttgtttttaatagataaaatatctttattttcttgatctgtttttattacagctattttattattattatcttta |
| 1b6403a067c0944bc7dc5d6463bedd72b075bd51 | Hymenoptera | Braconidae | Apanteles | Apanteles parasitellae | attatctttaattttaggtcatggtggaatatcagttgatataggtattttttctttacatttagctggtatttcttcaattataggagctattaattttatttcaacaatttttaatttacgtacaagtttatttgatatagataaaatatctttattttcttgatcagtttttattactgcaattttattattgttatcatta |
| a9de44228382821ba661fb8e90b708d5632290f4 | Hymenoptera | Braconidae | Aphidius | Aphidius cingulatus | tttatctttaactttaggtcatagaggagttgcagtagattttgctattttttctttacatttagctggtatttcatcaattataggagcaattaattttattagtactatttttaatatacgtccttataatattaaaatagatcaaatttctttattagtttgatcagttttaattacagctgttttattattattatcttta |
| e1c263b52411b55597c016aab8084971b914ebd5 | Hymenoptera | Braconidae | Aphidius | Aphidius eadyi | tttatcattaacgttaggacatagaggagcagcagtagattttgctattttttctttacatttagcaggtatttcttcaattataggtgctattaattttattagaactatttttaatatacgatgttataatattaaaatagatcaagtttcattattgatttgatcagttttaattactgctattttattattattatcatta |
| cf8fecb07c74b19559fc2a6793dab8a1b7a04ce4 | Hymenoptera | Braconidae | Aphidius | Aphidius ribis | cttatctttaactttaggtcataggggggttgctgtagattttgctattttttctttgcatttagctggtatttcttcaattataggagctattaattttattagaactatttttaatatacgatgtaataatattaaaatagatcaaatttcattattaatttgatcagttttaattactgctgtgttattattattgtcactt |
| c3ebe70d5539f694b1c728eec8e9f4e65285bfdf | Hymenoptera | Braconidae | Aphidius | Aphidius schimitscheki | attatccttaactttaggtcacagaggtgtagcagtagattttgctattttttctttacatttagcaggtatttcttcaattataggtgctattaattttattagaactatttttaatatacgatgttacaatattaaaatagatcaaatttcattattaatttgatcagttttaattactgctgttttattattattatcttta |
| a69531297d78935cb7a26ad6efc99a0e1a823e21 | Hymenoptera | Braconidae | Aphidius | Aphidius uzbekistanicus | tttatcattgactttaggtcatagaggtgtagcagtagattttgctattttttcattacatttagcaggaatttcttctattataggggctattaattttattagtacaatttttaatatacgatcttataacattaaaatagatcaaatttcattattggtatgatcagttttaattactgctgttttattattattatcatta |
| 4255f397ee18c4b510cb6961b39b83a1c93fcd7e | Hymenoptera | Braconidae | Ascogaster | Ascogaster provancheri | tttatctttaattattggtcatggtggtatttcagtagatttaagaattttttctttacatttagctgggatatcttcaattataggtgcaattaattttattgttactattataaatatatgatttggtttaaaatatatagataaaattagattatttacttgatcagtaataattacagcaattttattattattatctttg |
| e4f0d8801d0d9e2bf01ed06c59c24d0215d201f3 | Hymenoptera | Braconidae | Asobara | Asobara rufescens | tttatcttcaagtattggtcatagaggaatttctgtggatttagctattttttcattacatttagctggtgcatcatctattataggggtaattaattttttaacaacaatttttaatataaaattttatataattaaattagatcaattaagattatttgtatgatcaattttaattactgctattttattattattatcatta |
| 8b0ae3d21cd3e8efc0b46f74e0197341b47437ee | Hymenoptera | Braconidae | Binodoxys | Binodoxys acalephae | attatcattgaatttaggtcatagtggtatttctgttgatttagctattttttctttacatttagcaggtatttcatcaattataggtgcaattaattttattagaactattttaaatatacgttcctatagagtatctatagatcagatttcattattagtttgatcagtattaattacagctattttattattattatcatta |
| 7575234dca904cf137bebbf8fcb4e9bd3a81aad2 | Hymenoptera | Braconidae | Binodoxys | Binodoxys brevicornis | attatctttaaacttaggtcataggggtattgcagtagatttagctattttttctttgcatttagctggtatttcatcgattataggtgcaattaattttattagtactattttaaatatacgtgcatataatgtttcaatagatcagattccattattggtttgatctgtgttgattactgctattttattattattatcgtta |
| 0c94da2cb55f8b602e3a96aadd19adad054b5dfb | Hymenoptera | Braconidae | Binodoxys | Binodoxys centaurae | attgtctttaaatttaggtcatagtggtgttgcagtagatttggctattttttctttacatttagctggtatttcttctattataggtgctattaattttattaggactattttaaatatacgttcttatagggtttctatggatcaaatttctttactagtttggtctgtgttaattacagctattttattattattgtcttta |
| 1716df57b3c70a71b3bbb5244529725a8a4e266d | Hymenoptera | Braconidae | Chelonus | Chelonus andrievskii | attgtctttattaataggtcatagaggtgtatcagttgatataagaattttttctttacatttggctgggatatcatcaattataggttcaattaattttattgttactattataaatacatgaataaaatttagttttatagataaatttcctttatttgtttgatcagtttttattacaactattttattgttattgtcttta |
| 7350ccae920dc7269d2ce7a61ef0db2b6c84ec21 | Hymenoptera | Braconidae | Choeras | Choeras suffolciensis | tttatctttaattttaggtcatggtggtatatctgttgatataggaattttttctttacatttagcaggaatttcatcaattataggtgctattaattttatttcaactattttaaatttgcgtacaaatttatttaatatagataaaatatcattattttcttggtcagtacttatcactgctattttattgttattatcttta |
| fc3fec5083402bdeadf7938524ee9dbdaa723ced | Hymenoptera | Braconidae | Chorebus | Chorebus anasellus | cttatcatctagaattggacatggagggatatctgttgatttagctattttttctttacatttagctggtgtttcttctattataggagcaattaattttattactacgatttttaatataaatttttttataattaaattagatcaattaagtttatttatttgatctattttaattacagctattttattattattatcttta |
| 0e77013efe92e86116e56d7f88d24beebafe14f8 | Hymenoptera | Braconidae | Chorebus | Chorebus longicornis | gttatcttcaataattggtcatggtgggatatccgttgatttagcaattttttcattacatttagctggtgcatcttcaattataggagctattaattttattactacaatttttaatataaatttttttataattaaattagatcaattaagtttatttatttgatcaattttaattactgctattttattattattatcatta |
| dd93465f451250051741c6c6d4ce1dd30d5dbee6 | Hymenoptera | Braconidae | Chorebus | Chorebus parvungula | attatcatctagaattggtcatggcgggatatctgttgatttagcaattttttctttacatttagcaggggtatcttctattataggagctattaattttattactacaatttttaatataagtttttatataattaagttagatcaattaagattatttgtgtgatcaattttaattacagctattttattattattatcttta |
| 7ad18c45eb37b121112004a1855acda4fe107e65 | Hymenoptera | Braconidae | Cotesia | Cotesia pilicornis | tttatctttaattttaggtcatggtggaatatcagttgatttaggaattttttctttacatttagctggtgcttcttcaattataggtgcagttaattttattactacaattataaatatacgttctaatttatttaatatagataaaatatctttattttcttgatcagtatttattactgcaattttattattattatcttta |
| 695441a8b5e266a90155befb58759639c859642a | Hymenoptera | Braconidae | Cotesia | Cotesia rubecula | tttatcattaattttaggtcatggtggaatatctgtagatttaggaattttttctttacatttagctggagcatcttctattataggtgctgtaaattttattactactattataaatatacgttcaaatttatttaatatagataaaatatctttattttcttgatctgtatttattactgcaattttattattattatcttta |
| d7fbca5abe631b9ce93b92237bb96feacd4f8331 | Hymenoptera | Braconidae | Cotesia | Cotesia tenebrosa | tttatctttaattttgggccatgggggaatatcagttgatttaggaattttttctttacatttagctggtgcttcttcaattataggtgctgttaattttatttctacaattataaatatacgttctaatttatttaatatagataaaatatccttattttcttgatcagtatttattactgcaattttattattattatcttta |
| da42908e9e4d66624c0e134ed473c845cc54b7d1 | Hymenoptera | Braconidae | Cotesia | Cotesia vanessae | tttatctttaattttaggacatagtggtttatctgttgatttaagaattttttctttacatttagctggtatatcttcaattataggagcagttaattttattacaacaattttgaatatacgttctggtatatttaatatagataaaatatctttattttcttgatcagtatttattactgcgattttattattattatcttta |
| 367f27268e31546262957a487bf0bdde4f176b65 | Hymenoptera | Braconidae | Cotesia | Cotesia xylina | tttatctttaattttaggtcatggtggtatatctgttgatttaagaattttttctttacatttagctggtgcatcttcaattataggtgctgtaaattttattacaacaattttaaatatacgaactaatttatttaatatagataaaatatctttattttcttgatcagtatttattactgcaattttattattattatcttta |
| c30406b75b343726499ea083c510242654526131 | Hymenoptera | Braconidae | Dacnusa | Dacnusa areolaris | tttatcttctataattggtcatggtggaatatctgttgatttagctattttttctttacatttagcgggggtttcttcaattataggagctattaattttattactacaattttaaacataaatttttttataattaaattagatcagttaagtttatttatttgatcaattttaattactgcaattttattattattatcttta |
| db4b6d3a6d99b9ed67010be5d34397551fd43cb2 | Hymenoptera | Braconidae | Diaeretellus | Diaeretellus svalbardicum | cttatctttaactttaggacatagaggtgtggcggtagattttgctattttttcactacatttagcaggtatttcttcaattataggagctattaattttattagtactatttttaatatacgatcatataatattaaaatagatcaaatttcattattaatttgatcagttttaattactgctattttattattattatcatta |
| fd62d3b9b5752d768710e3d5424becf68452de13 | Hymenoptera | Braconidae | Distatrix | Distatrix formosus | attatctttaattattggtcatagaggtatatcagttgacataagaatcttttctcttcatttagcaggtgcatcttcaattataggagcaattaattttatttcaacaatttttaatatacgtacttttttttttgaaatagataaaatttcattattttcttgatctgtattaattaccacaattcttttacttttatcatta |
| ae68b3efcc2c2b354bdd821cf35aa039d272a377 | Hymenoptera | Braconidae | Dolichogenidea | Dolichogenidea lineipes | attatctttaattttaggacatggtggtatatcagttgatttaggaattttttctttacatttagcaggtgcttcatcaattataggagctgttaattttattactactattttgaatatacgtacaaatttatttttaatagataaaatgtctttattttcttgatctgtttttattacggcaattttattattattatcatta |
| 8b595436e5845508acf3d1cb1c139a981432e59d | Hymenoptera | Braconidae | Dolichogenidea | Dolichogenidea phaloniae | tttatcattaattttaggacatggtggtatatcagttgatttaggtattttttcattacatttagctggtgcttcatcaattataggtgctgttaattttattacaacaattttaaatatacgaacaaatttatttataatagataaaatatctttattttcttgatcagtttttattactgcaattttattattattatcatta |
| 51801aa69a3d56acda86428c0ef072458afa1da3 | Hymenoptera | Braconidae | Ephedrus | Ephedrus koponeni | attatcattaaatttaggacatagaggtatatcagttgatatctctattttttctttacatttagctggtatttcttcaattatgggggctattaattttattacaacaattttaacaataaattctttaggaatagttaaagatcaattacctttattatgttgatcaattattattacagcaattttattattattatcttta |
| 3ecadcb0c48b12ee6bfeefeffb44443420218e20 | Hymenoptera | Braconidae | Ephedrus | Ephedrus nacheri | attatcattaaatttaggacatagaggtatatcagttgatatttctattttttctttacatttagctggtatttcttcaattataggggctattaattttattacaacaattttaacaataaattctttagggatagttaaagatcaattacctttattatgttgatcaattattattacagcaattttgttattattatcttta |
| 36355b2e0498956e193138c2c814ebc09076cf2a | Hymenoptera | Braconidae | Ephedrus | Ephedrus niger | attgtctttaaatttaggtcatagtgggatagctgttgatatttcaattttttctttacatttagctggtatttcttctattataggagctattaattttattacaacaattttaacaataaatcctttaggaatagttaaagatcaattgccattattatgttgatcaattattattacagcaattttattattattatcttta |
| 09bf1a195946b735277c6798143acd524b573bdb | Hymenoptera | Braconidae | Habrobracon | Habrobracon crassicornis | tttatcatcttctttaggacatagaggtttatctgttgatttagctattttttctttacatattgctggaatttcatcaattttaggggctattaattttattactactatttttaatatacatttatttattttaaaattagatcaattaactttattaatttggtcaatttttattactgctgtattattattattatcttta |
| 6a05d14554a226bfc3993424c0d75554eb1b762b | Hymenoptera | Braconidae | Habrobracon | Habrobracon hebetor | tttatcttcttctttaggtcataacggagtatcaatagatttaacaattttttctttacatttagcaggaatttcttctattttaggatcaattaattttatttctacaattttaaatatacatttatttactttaaaattagatcaattaactttattaatttgatcaatttttattactacaattttattattattatcctta |
| 75e6b4d4d48a60cfc4d907d5f3d4a12993140c5e | Hymenoptera | Braconidae | Leiophron | Leiophron duploclaviventris | cttatctttaaatttaagtcatgcaggaatatctgtagatttagctattttttctttacatttagcaggaatatcttcaattataggggcaattaattttattactactattattaatatacgtttattagggttaaaaatagataatatttctttatttacatgatcagttaatattacagcaattttattgttattatcttta |
| d5e7af08a03334d5ee25366eec94b1bf4f7db780 | Hymenoptera | Braconidae | Leiophron | Leiophron similis | attatcattaaatttaagtcattctaggatatctgttgatttagcaattttttctttgcatttagctggtatatcatcaattataggagctattaattttattactactattattaatatacgtttatgagggttaaaattagataatattactttattttcatgatctgttaatattacagctttattattattattatcttta |
| dec9f21e30b22fda740461ba7de6de94fe7cf148 | Hymenoptera | Braconidae | Microctonus | Microctonus aethiopoides | tttatctttaaatgtaagtcataggggaatatctgttgatataagaattttttctttacatttagcagggatttcttctattataggagcaattaattttatttctactattataaatatacgtttaataggtttaataataaataatatttctttatttgtatgatctgtgttaattactgctgtattattattattatcatta |
| e18e8e002aa83719a3156770896945de1ee76a97 | Hymenoptera | Braconidae | Microplitis | Microplitis deprimator | tttatctttaattttaggtcatggtggtttatctgtagatttaagaattttttctttacatttagctggtgtttcttcaattataggagctgtaaattttattacaacagtttataatatgcgttctaattttttaaatatagataaaatttctttatttatttgatctgtattaattacagcaattttattattattatcttta |
| bb3d0f50388e93b2b61a42586c4a249198555d96 | Hymenoptera | Braconidae | Opius | Opius ocreatus | attatcatctatggttggtcatggtggtttatcagttgatttagctattttttctttacatttagctggtgtttcttcaatcataggagctattaattttattacaactatttttaatataaatttttatataattaaattagatcaattaagtttattaatttgatcaattttaattacggcaattctattattattatcatta |
| 3097096e02fd2a69674e8fb16b88a073a56eeed2 | Hymenoptera | Braconidae | Praon | Praon barbatum | attatctttaattaatagacatagaggaatttcagtagatttagcaattttttcattacatttggctggaatttcttcaattataggagcaattaattttatttcaacaattttaaatatacgattaaatgatataactatagatcaaattcctttatttgtttgatcagtttttattacagttattttattacttttatcttta |
| 80db6465de78b4b42a1980bdbbc26adf2aec066d | Hymenoptera | Braconidae | Praon | Praon yomenae | tttatctttaattaatagacatagaggaatttcagtagatttagcaattttttcattacatttagctggtatttcttcaattataggatctattaattttatttcaacaattttaaatatacgattaaatggtataactatagatcaaattcctttatttgtttgatcagtttttattacagtaattttattactattatcttta |
| c19882fbf13eced9b80233949ed1189fbb37a851 | Hymenoptera | Braconidae | Protapanteles | Protapanteles porthetriae | tttatcattaattttaggtcatagaggaatatcagttgatatgggaattttttctttgcatttagctggtgcttcttcaattataggtgcagtaaattttattactacaattttaaatatacgaacaaatttatttaaaatagataaaatatctttattttcttgatctgtatttattactgctattttattattattatcttta |
| 6d06367c95f0bd114acb8edbedde4f1370840c9b | Hymenoptera | Braconidae | Remaudierea | Remaudierea plocamaphidis | tttatcattaactttaggtcatagaggggttgcagtagattttgctattttttctttacatttagctgggatctcatcaattataggagcaattaattttattagtactatttttaatatacgttcttataatattaaaatagatcaaatttctttattagtttgatcagttttaattacagctgttttattattattgtcttta |
| a87c1a54fa3e74406ab0b2da7d3949d40bf11758 | Hymenoptera | Braconidae | Sathon | Sathon lateralis | tttatctttaattttaggtcatagtggtatatcagttgatatgggtattttttctttacatttagctggtgcttcttcaattataggagcagtgaattttattactacaattttaaatatacgaacaaatttatttaaaatggataaaatatctttattttcttgatctgtttttattactgctattttattattattatcttta |
| 5829c1873f770c91add6d72fa3d983f4c881bc81 | Hymenoptera | Braconidae | Triaspis | Triaspis pallipes | attatctttaaatattggtcatggaggaatatcagttgatttagcaattttttctttacatttagcaggaatttcatcaattataggagctattaattttattacaactattttaaatatacgtcctaaaataattactatagataaaatttctttacttagatgatctattttaattacagctattttattattattatcatta |
| 64db68f9f16f157fa9d9ba8ddcbd2416fd4be712 | Hymenoptera | Braconidae | Trioxys | Trioxys pallidus | tttatctctaaatttgggacatagtggtatttcagttgatttagctattttttctttacatttggcaggtatttcttcaattatgggggctattaattttattagaacaattttaaatatacggccatgtggagtttctatagatcagattcccttattagtttgatctgttttaattactgctgttttgttactattatctttg |
| c9f21930a0d61cbfb9bd1dcf4e46b1f0dae46309 | Hymenoptera | Braconidae | Trioxys | Trioxys sunnysidensis | attatctttaaatttaggacatagaggtatttctgtagatttagctattttttctttacatttagcaggaatttcttcaattataggggcaattaattttattagaactattttaaatatacgttcttatagagtttcaatagatcaaatttctttattagtttgatctgtattaattacagcaattttattattattgtcttta |
| 055138c17022d394e2eeeec35fc1f964c8cd99b2 | Hymenoptera | Charipidae | Alloxysta | Alloxysta pleuralis | tttatcttctaatttaggccatgcaggtatctcagtggatttaacaattttttctttacatataagaggaatctcatctattttaggttcaattaattttattacaactattttaaatatacgtcctattaatttatctatagataaaatttcgttattttcttgatcaattttattaacaactattttattattattatcttta |
| c1e9da60eef271e91a47ec99bba7ae4e1873c54e | Hymenoptera | Crabronidae | Miscophus | Miscophus bicolor | tttatcatcattattaggtcataatgattgttctgttgatatagctatttttgctttacatattgggggtgcttcatctattataggttcaattaattttattgttactattattataataaagaataaaagtttaagtatagatcaacttcctctttttatttggtctgttttgattactacaattttattattattatcttta |
| f797fcb9e31cc4e6f94969b970711b31e7e7150d | Hymenoptera | Crabronidae | Spilomena | Spilomena beata | tttatcatcaattagatttcataatagtccctccgtagatataactattttctctttacatattgcaggaatatcatctattataggagcaattaattttattgtaactattataaatttaaaaaataataatataacaatagatcaaattccattatttgcatgatcagtattaattactgctattttactattattatcctta |
| 73d9eea7f636d327458bddee60a70d99b046aaeb | Hymenoptera | Cynipidae | Aulacidea | Aulacidea subterminalis | cttatcttcaaatataggacacatagggatttcagttgatttaacaatttattctttacatataagagggatttcttcaattttaggttcaattaattttattacaacaattttaaatatacgccctaaaataatatctatagataaaatttcattatttatatgatcaatttttcttacaacaattttattattattatcttta |
| ab0a89ec554a195d10659910eff0f2f5a0b94f74 | Hymenoptera | Diapriidae | Monelata | Monelata solida | tttatctacaaatatttttcataatgatatatcagtagattttactattttttcattacatattgcaggtatttcttctattataggagcaattaattttattagaacaattattaatatacgatcaaaattaataaatttaaatttaattagattattttcatgatcaatttttttaacagtaattttattattattagcatta |
| 7245bd6d60534022302e0ee6f795a0aa2d9e2fc2 | Hymenoptera | Dryinidae | Anteon | Anteon tenuicorne | cctatcaagtaatatttctcatagaagtatatctgttgatttaactattttatctctacacttagctggggcctcatctgttataggggcaattaatttcattgctactattataaatataacttgttttaatattaaattaactcaaattaatttatttatttgatcaatttttattactgtaattttattattattatcatta |
| c9605c518e0fda7076ce3f2ee36792074e2c88f4 | Hymenoptera | Figitidae | Kleidotoma | Kleidotoma filicornis | tttatccttaagtgcaagtcacccaggtatttctactgatttagtaatttactcccttcatttaagaggagtatcttctattttaggatcaattaatttcatttcaactattttaaatgtacgtcctaactatatacttatagataaaatttctttatttgcctgatcaatttttttaacaacaattctccttttactctcttta |
| 18c7a6481690471358232a6c03645c187652e835 | Hymenoptera | Ichneumonidae | Atractodes | Atractodes cryptobius | acttgcaaataatattaatcatgaaggaatatcattggattttgctattttttcattacatatagctggtttatcttcaattataggtgcaattaattttatctcaacaattataaacatacgaacaattaatataacctttgataaaatttctttattttgttgatcaattaaaattactgttattttacttttattagcagtc |
| b7e0f00eed446a0994a0d15cf736d94a4dc073d7 | Hymenoptera | Ichneumonidae | Atractodes | Atractodes fumatus | cctctcattaaatattaaccacgaaggtatatcaattgatatagctattttctctctacatcttgctggtatatcatcaattataggagcaattaattttattacaactattataaatatatttccattaaaattaaaatttgaacaattaactttattcacatgatcaattttaattacaacaattttattattaattgcagtt |
| a164356c2a6621d6d2138556b989437fb0c91542 | Hymenoptera | Ichneumonidae | Barichneumon | Barichneumon plagiarius | tttatcatctaatttaaatcatgaaggtttatcagtagatatatcaattttttctcttcatttaactggaatatcatcaattataggagccattaattttatttcaacaattttcaacatatatcctattaatttaaaatttgaacaattaacactatttacttgatcaattataattacaacaattttacttcttttagctgta |
| ad992b1843ae706e43fa6d7956864c1605e1b193 | Hymenoptera | Ichneumonidae | Campoletis | Campoletis latrator | attatctttaaatattaatcatgaaggtatatcaattgatttatcaattttttcacttcatttagcaggaatatcttcaattataggagcaattaattttattactacaatttttaatataaaaaatattaataaaaaatttgaacaattaactctatttacttgatcaattaaaattactactattttattacttttagcagtt |
| 8e40ef54fa2c2dbd5a42fbf49a276a3abbfabeb5 | Hymenoptera | Ichneumonidae | Campoletis | Campoletis thomsoni | ctatccttaaatattaatcatgaaggaatatcagtagatttatctattttttctttacaccttgcaggtatatcatcaattataggagcaattaattttatctcaacaattattaatataaaaaatattaataaaaaatttgaacaattaactctatttacttgatcaattaaaattacaactattttattattattagcagtc |
| ed6912ae5c797babb4ed82744254c9b1fa9ae309 | Hymenoptera | Ichneumonidae | Demopheles | Demopheles corruptor | tctttctcttaatattaatcatgaaggaatatctgttgatatagcaattttttctattcatttagctggtatatcatcaattataggggcgattaattttatttctactattataaatataaaaatttcaaattcaaaatttgaacaattaactctattttcatgatcaattattattacaacaattttattattattagctgta |
| ccb360ef951159f3c973b4511a48038e16d01166 | Hymenoptera | Ichneumonidae | Diadegma | Diadegma erucator | tttatcattaaacataagacatgaaggaatatcagtagatttagctattttctccttacatcttgcaggaatatcatcaattataggagcaattaattttattacaactatttttaatataaaaaattataataaaaattttgaacaactcactttatttacctgatcaattaatattacaacaattttacttttattagcagtt |
| a546345744e852e4849bb4c7ffe50717a6c401ed | Hymenoptera | Ichneumonidae | Diadegma | Diadegma semiclausum | actttcattaaatattagacatgaaggaatatctgtagacttagctattttttcattacatcttgcaggaatatcatcaattataggagcaattaattttattacaactatttttaatataaaaaattataataaaaattttgaacaattaactttatttacttgatcaattaatattactacaattttacttttattagctgtt |
| 3a979df4f954e177ab667b55d64514e52ee33e5f | Hymenoptera | Ichneumonidae | Enytus | Enytus neoapostata | cctatccttaaatattagtcatgaaggaatatctattgacttagcaattttctctctacacttagcaggaatatcctcaattataggagcaattaactttatcacaactatttttaatataaaaaatttaaaaaaaaattgagaacaattaactttatttacatgatcaattaatatcacaacaattcttcttttattagcagtt |
| 7bc121e6bc3d86101f28d64a290d0d3f8eb08928 | Hymenoptera | Ichneumonidae | Eridolius | Eridolius dahlbomi | tttatcacttaatttaagtcatgaaggtatatcagtagatttaacaattttttctcttcatttagccggtatatcttccattataggagcaattaattttattacaacaatttttaatatacgatcaaaaaatataatattagaacaaatatcattatttacatgatctattaaattaactacaattcttttattactagcagtt |
| 47813d3cc87b6410cce2b7b0ae661f7f288e9143 | Hymenoptera | Ichneumonidae | Heterischnus | Heterischnus pulex | cttatctttaaatttaaatcatgaaggaaattctgttgatatagcaattttttctcttcatatagctggtatatcttcaattataggagcaattaattttattacaactatccttaatataaaaattttaggatcaactcttgaacaaataactttatttgcatgatcaattcaaatcacagctattcttcttttattagctgtt |
| 126d4bdcf62a21300794f8cf3b6a567c397efa51 | Hymenoptera | Ichneumonidae | Hypsicera | Hypsicera ecarinata | cctttctctcaatattagtcatgaaagtatatctgttgatttcagaattttttcccttcatcttgcaggtatatcttctattataggagccattaatttcattacaactattataaatataaaaatctctaatatttcatttgaacaaataactttattctcctgatctattcaaattacagcaattttattattattagctgtt |
| c13595185fc3fb079a79dfbed8495112c4aa0cb3 | Hymenoptera | Ichneumonidae | Lysibia | Lysibia nana | attatcattaaatttaaataatgaaggaatatctattgatatttcaattttttcattacatttagcaggaatttcatcaattataggtgcaataaattttattagatcaattataaatataaatccaaataatttaaaatttgaacaattaagattattttcatgatctattttaattacaacaattcttttattattagcagta |
| 0d55a8b54562dfdb1ec73fa2aa3154ce73568a39 | Hymenoptera | Ichneumonidae | Mastrus | Mastrus ridibundus | tctatctttaaatattaatcatgaaggaatatcagtagacttatctattttctctttacatcttgctggaatatcatcaattataggagctattaattttattactactattttaaatatttatcctataataataaaaattgaacaattaactttatttacatgatcaattttaatcacaactattttacttttattagctgtc |
| d65e6e7e3ba179f469ebbc3e476ceb4b6e7c308b | Hymenoptera | Ichneumonidae | Mesoleptus | Mesoleptus laticinctus | cctatcattaaatttaaatcatgaaggcttatccattgatatagcaattttttctttacatttagcaggtatatcctcaattataggagctattaattttattactactatcataaatatatttcctattataataaaatttgaacaattaactcttttttcttgatcaattttaattactacaattttacttttacttgcagtc |
| 3002973a0497e93ab89eeb5598cc606e30616b87 | Hymenoptera | Ichneumonidae | Micromonodon | Micromonodon tener | cttatccttaaatttaaatcatgaaggtatatcagttgatttatctattttttctcttcatttagcagggatatcctctattataggagctattaattttattagaactatcataaatatattcccaacaaatataaaatttgaacaactaacattatttacatgatcaattttaattaccaccattcttttattactagcagta |
| 726f707f407c79f1f775f50554366341bb00d16f | Hymenoptera | Ichneumonidae | Neurateles | Neurateles falcatus | cctatccctaaatattaatcatgaaggaatatcagtagatatatcaattttctccttacatttagctggaatatcatcaattataggagctattaactttattacaacaattttaaatatacgacctaattcaattaaattagacaaaatctcattatttacatgatcaattaaaattactacaattttactattattggctgta |
| c522cec92e0a4147367580c484d673487dc49729 | Hymenoptera | Ichneumonidae | Orthizema | Orthizema gravipes | actatcactaaatttaagacatgaaggtgtttcaattgacatagcaattttttctattcatttagcaggaatatcctcaattataggagcaattaattttattactactattataaatatatttcccttaaatataaaattagatcaacttactttatttacatgatctattttaattacaacaattttattattattagcagtc |
| 7cbefa8017c8515fabcdec12863bc8eaa4170bce | Hymenoptera | Ichneumonidae | Phrudus | Phrudus monilicornis | cttatctttaaatctaaatcatgaaggaatatcagttgatgtatctattttttccctacacttagcaggagcatcttcaattataggagctattaattttattaccactattctaaatatacgtattataggaattctattgaatcaaataccattatttatttggtccattaaaattacagctattcttttattattagcagta |
| c51ccffb78fbd802b1bfafb007865472d7b94711 | Hymenoptera | Ichneumonidae | Rhyssa | Rhyssa amoena | cttatcccttaatattaaccatgaaggattatcagtagacttagcaattttctcattacatatagcaggaatatcctctattataggtgcaattaactttatttcaacaattattaatatacgacctcatataattaatatagaaaaaatatctttattttcttgatcaattataattactgcagtacttttacttttagcagta |
| 8240ae4301d8579726dd13513ca51fbd5b917dc2 | Hymenoptera | Ichneumonidae | Scambus | Scambus hispae | cttatcattaaatttaaatcatgaagggttatcagttgatttagcaattttttcccttcatatggcaggtatatcatctattataggagctattaattttattacaactatcataaatatacgacctaatataattacattagaaaaaatatctttatttacttgatcaattaaaattactgctattttacttttattagccgtc |
| e3e497377385345370a0d978adea1050b9b1444c | Hymenoptera | Ichneumonidae | Stenomacrus | Stenomacrus affinitor | cctatctttaaattaccatccagggatatctattgatatagcaatcttttcattacatttagcaggaatatcatcaattataggagccattaattttattaccacaattttaaatatacgacctaattcaattaaattagataaaatttctttatttacttgatcaattaaaatcactacaattttacttttattagctgtt |
| 0c681613ce421b1e591cdf20fa394a566bbe97d8 | Hymenoptera | Ichneumonidae | Stenomacrus | Stenomacrus deletus | tctttcattaaatatcaatcacgaaggaatttgtgttgatttagcaattttttctttacatttagctggtatatcttcaattataggagcaattaattttattactacaatttttaatatacgaccaaaattaattagtttagataaaatttcactatttacatgatcaattttaattaccactattttactattactggcagtt |
| e8a5e7cfb232df0dc7ffcc3a52ceeb419c4a0d3b | Hymenoptera | Ichneumonidae | Symplecis | Symplecis leucostoma | tctctcattaaatgtaaatcatgaaggaatagcactagattttgcaattttttccctccacatagctggcatatcatcaattataggagcaattaattttattacaacaattattaatatacgaactaaatttataagattggataaaatatctttatttatttgatcaatcaaaattactgcaattctattattactcgcagta |
| da70ce04e80135ebd7063f28498213566351d8bd | Hymenoptera | Ichneumonidae | Woldstedtius | Woldstedtius citropectoralis | tttatcatctaatttaaatcatgaaggaatatcattagatttagctattttttctttacatttagccggaatatcttctattataggagcaattaactttattactacaattattaatatacgacctaataatttaaatttagaaaaaatatcgttattttcatgatctattttaattacagcaattttattattattagctgta |
| 4dc72fb85cf1f2f202e46d63d2dce9d51275bd9c | Hymenoptera | Tenthredinidae | Cladius | Cladius pallipes | tttatcaagaagaatttcacatgcaggagcatctgtagatttaactattttttcattacatttagctggaatttcatcaattcttggggctattaacttcatctcaacaataattaatatacgagtaaaaggaataaatttcgaacgaatacctttatttgtatgagcagtatcactaacagctttattacttcttctatcccta |
| 298d60192a6518581a1b3f54b2f314b2b7666f4d | Hymenoptera | Tenthredinidae | Cladius | Cladius pilicornis | tttatctaatagaatttcccatgcaggggcatctgttgatctaacaattttttctcttcatttagcaggaatttcatcaatccttggagctattaattttatttcaacaataattaatatacgagtaaaaggaataagatttgaacgaatacctctatttgtttgagcagtatcacttacagcattattacttctactatcatta |
| 1bcfcb4d4845ba7dfe9270eeb678e3aea3ffcc3c | Hymenoptera | Tenthredinidae | Nematus | Nematus bergmanni | attatcaagaagaatctctcattcaggggcatctgtagacttaacaattttttcattacacttagcaggaatttcatcaattttaggggcaattaattttatttcaacaataattaatataaaactaaaaggaataagatttgaacaaatacctttatttgtatgagcagtttcattaacagcattattactactattatcacta |
| d3317123ca9236a8e9a4296316d416edbe021f36 | Hymenoptera | Tenthredinidae | Pristiphora | Pristiphora albitibia | cctatcaagaagaatttcgcatgcaggggcatcagttgatttaactattttttcattacatttagcaggtatttcatcaattttaggggcaattaattttatttcaacaataatcaatataaaacttaaaggaataaaatttgaacaaatacccctatttatttgagcagtatcattaacagccttattacttcttttatcatta |
| a1ba9bdab07a54bf2a516dd53837794db2b56502 | Psocoptera | Ectopsocidae | Ectopsocus | Ectopsocus meridionalis | cctatcaagtgttattgcccatacaggggcatctgttgatttagctattttttcacttcatttagctggtgttagttcaattttaggtgctgtaaattttattacaacaattattaatatacgatcaaacggtttaactttagaacgtatacctttatttgtttgggctgtatttattacagcaattttattattattatctctt |
| 63e137c75dea6f47b4bfab5c5f2305b38fccb2ec | Psocoptera | Liposcelididae | Liposcelis | Liposcelis rufa | ctatcattatattcagcccatcccagtgaaagtgtagacttagcaattttttctcttcacttagctggagcgagatcgattttaggagctattaattttattactacattttttaatctatgagtttttaaaaataaaatagaattgagaagtttatattcttggtctgtttcaatcacagcagttttacttttactttcttta |
