## Supplementary material for "Detecting flying insects using mega-nets and meta-barcoding": Table_S4

Table S4 We detected 15 ASVs present in 25% of all the car net samples (corresponding to <7% of all sequences) belonging to eight named species and six ASVs that could not be assigned to species level. These are arguably the most common airborne animals in Denmark (at least in June).

| **Species** | **Biology & notes** |
| --- | --- |
| *Brassicogethes aeneus* | pollen beetle |
| *Schizohelea leucopeza* | biting midge with two sequence variants |
| *Smittia albipennis* | non-biting midge |
| *Empis aestiva* | predatory fly |
| *Sciara atomaria* | fungus gnat |
| *Metopina oligoneura* | generalist scuttle fly |
| *Rhopalosiphum padi* | cereal crop pest |
| *Delia florilega* | leaf miner |
| *Rachispoda* sp. | dung fly |
| Scatopsidae sp. | dung midge |
| Thysanoptera sp. | thrip |
| *Smittia* sp. | non-biting midge |
| *Oscinella* sp. | grass fly |
| *Corynoneura* sp. | non-biting midge |
