## Supplementary material for "Detecting flying insects using mega-nets and meta-barcoding": Table_S3

Table S3 The 29 species with no occurrence records for Denmark, Sweden, Norway and Germany in GBIF and www.allearter.dk and associated sequences used for taxonomic assignment. Further comparison with the Fauna Europea database (<https://fauna-eu.org/>) and the BOLD taxonomy database (<https://www.boldsystems.org/index.php/TaxBrowser_Home>) revealed that 11 species did have occurrence data for Sweden, Norway and/or Germany based on the two other databases.

| **ASVID** | **order** | **family** | **genus** | **species** | **Fauna Europea DK occurrence** | **Fauna Europea Sweden, Norway, Germany occurrence** | **BOLD taxonomy DK occurrence** | **BOLD taxonomy Sweden, Norway, Germany occurrence** | **Occurrence based on GBIF, BOLD, Fauna Europea** | **Biology (if known) & comments** | **sequence** |
| --- | --- | --- | --- | --- | --- | --- | --- | --- | --- | --- | --- |
| e57672d1054792f02888bba2c7c87d0dccda3c03 | Diptera | Agromyzidae | Amauromyza | Amauromyza karli | no | no | no | no | Canada, China, Bulgaria, Belarus, Russia, Poland, Finland | Leaf miner | attatcttcaattattgctcatggaggagcttcagttgatttagctattttttctttacatttagccggagtttcttcaattttaggggcagtaaattttattactacaattattaatatacgaactactggaattacatttgatcgaatgcccctttttgtatgatctgtattaattacagctgtattattattactttcttta |
| 6a0dfc414f34d2a2da5b9c54ab31d05df85a9c98 | Diptera | Anthomyiidae | Botanophila | Botanophila relativa | no | Sweden, Norway | no | no | Finland, Nearctic region, Norway, Sweden, United States of America, Switzerland, Canada |  | tttatcttctaatattgctcatggtggagcttctgttgatttagctattttttctttacatttagcaggaatctcatctattttaggagctgtaaattttattacaactgtaattaatatacgatcaacaggaattacttttgatcgaataccattatttgtatgatctgtagtaattacagctttgttacttttattatcttta |
| d39626e7601bcb3c1224f0d93931b7797c1539b1 | Diptera | Chironomidae | Tanytarsus | Tanytarsus excavatus | no | Sweden, Germany | no | no | Found in 14 countries: <https://fauna-eu.org/cdm_dataportal/taxon/3e331460-a99b-4ef9-acba-b7ea072d5000> |  | cctttctgctaatattgcccatagaggagcatccgtagatcttgctattttttctttacatttagcaggaatttcttcaattttaggctcagtaaattttattactacagctattaatatacgagctaatggaatcactctagatcgaatacctttatttgtctgatctgtaattattacaacaattttacttcttctttctcta |
| 91034c07c6982ab28ea57dc88a34dc68da2e7f2d | Diptera | Dolichopodidae | Campsicnemus | Campsicnemus alexanderi | no | no | no | no | Canada, United States of America |  | tttatctgcaggaattgctcacggaggtgcttctgttgatttagctattttctctcttcatttagccggaatttcttctattctaggagctgtaaattttattacaacagtaattaatatacgatcaacaggaattacttttgatcgtataccattatttgtatgatctgttgttattactgctattttattattactttcatta |
| c9ede0d235793ac474f8961c19ea85455e3a41d5 | Diptera | Dolichopodidae | Dolichopus | Dolichopus celeripes | no | no | no | no | Canada, United States of America, Finland |  | cttatcctcaggaattgctcatggaggagcatctgtagatttagcaattttttcattacatttagcaggtatttcctcaattttaggggcagtaaattttattacaacagttattaatatacgatctacaggaattacatttgatcgaatacccctttttgtatgatcagtagtaattactgctattttattgttattatccctc |
| cb87397e15d94e62728efa271446b96bce38c213 | Diptera | Pipunculidae | Cephalops | Cephalops hardyi | no | no | no | no | Canada, United States of America, Russia | Parasitoid (family Pipunculidae) | tttatcttctaatattgctcatggtggagcctctgtagatttagctattttttcattacatctagctggaatttcatctattttaggagcagtaaattttattacaactgtaattaatatacggtcaaaaggaatatcttttgaccgaataccattatttgtgtgagctgtagctattacagctttattattattattatcttta |
| cb8a9187ed245dbde4634210fccfa6f63b8519bc | Diptera | Sciaridae | Bradysia | Bradysia procera | no | Germany | no | no | South Korea, China, Russia, France | Greenhouse pest (genus Bradysia) | tttatcttcaacaatttctcattcaggagcatcagtagatttatcaattttttcattacacttagcaggaatttcttctattttaggggcagtaaattttatttctacaattattaatataaaggcacccggtttatcatttgataaaattcctttatttgtttgatctgttttaattacagctattttacttttattatcatta |
| 5340c19623045b012cd5450347df0742b2f46367 | Diptera | Sciaridae | Camptochaeta | Camptochaeta cursor | no | no | not found | not found | Canada |  | attatcatcaacactatctcattctggtgcttcagtagatttatctattttttctctacatttggctggtatttcttcaattttaggagctgtaaattttatctctactatcattaatatacgaacccctggtatatcatttgataaaataccattatttgtttgatcagtattaattacagccgtattgttgttattatcttta |
| 1492cb593bbf9cc86a10317d8c3f198d75e1a1e4 | Diptera | Sphaeroceridae | Ischiolepta | Ischiolepta intermedia | not found | not found | no | no | Canada, United States of America |  | tttatcagcaggaattgctcatggaggagcatctgttgatttagctattttttcattacatttagctggaatttcttcaattttaggagctgtaaattttatcacgactgttattaatatacgatcaactggaattacatttgaccgaatacctttatttgtttgatctgttgtaattactgctttattattacttttatcatta |
| d7cef18f5f2b8f368cb52f92d9b2c4ce931f2553 | Diptera | Syrphidae | Parasyrphus | Parasyrphus vockerothi | not found | not found | no | no | Canada |  | tctttctgctagaattgctcatggaggagcttctgttgatttagctattttttctcttcatttagctggtatatcttcaattttaggagcagtaaattttattactacagtaattaatatacgatctaatggactttcttatgatcgaatacctttatttgtatgatcagtagtaattacagctttattattacttttatcatta |
| 9f9a4fc41f82e850fd3fd8fec10686dca6b6fb50 | Hemiptera | Aphididae | Aphis | Aphis neospiraeae | not found | not found | no | no | Russia, South Korea |  | tttatcaaataatattgctcataataatatttcagttgatttaactattttttctcttcatttagcaggtatttcatcaattttaggagcaattaattttatttgtactattttaaatataataccaaataacataaaattaaatcaaattcctctatttccatgatcaatcttaattacagctatactattaattttatcttta |
| 1678c65340d044da597f68fad3ad727cb64fac12 | Hemiptera | Aphididae | Hyadaphis | Hyadaphis passerinii | no | Germany | no | Germany | 21 countries:  <https://fauna-eu.org/cdm_dataportal/taxon/ebb3a07a-2d4b-49b4-8658-f35771fcd06f> & Canada, United States of America, New Zealand | Pest on Caprifoliaceae and Apiaceae | attatcaaataatattgcacataataatatttcagtagatttaactattttttctttacacttagcaggaatttcatcaattttaggagcaattaattttatttgcacaattcttaatataatacctaataatataaaaattaatcaaattcctcttttcccatgatcaatcctaattacagctattttattaattttatctcta |
| 045c2c5e2597539927e86fd3dbc9d6e72bac8353 | Hemiptera | Aphididae | Lipaphis | Lipaphis pseudobrassicae | not found | not found | no | no | 12 countries: <https://www.boldsystems.org/index.php/Taxbrowser_Taxonpage?taxon=Lipaphis+pseudobrassicae&searchTax=Search+Taxonomy> |  | tttatcaaataatattgctcacaataatatttcagttgatttaactattttttctctacacttagcaggaatctcttcaattttaggagcaattaattttatctgcacaatcctaaatataatacctaataatataaaattaaatcaaattcctcttttcccatgatcaattttaattacagctattttattaattttatctcta |
| 0b773750c895c330693e9f74e3f1f9aff27f4a84 | Hemiptera | Aphididae | Tuberculatus | Tuberculatus remaudierei | not found | not found | no | no | Belarus, Bulgaria, Canada | May be widely distributed in Europe, also found in Spain (<https://agris.fao.org/agris-search/search.do?recordID=US201302810483> and <https://ddd.uab.cat/pub/orsis/02134039v9/02134039v9p85.pdf>) and Greece (<https://www.researchgate.net/publication/234189298_A_contribution_to_the_aphid_fauna_of_Greece>) | actatccaataatattgctcataataatatctcagtagatttaactatcttttccttacacttagcaggaatctcatcaattttaggagcaattaattttatttgtacaattcttaatataatacctaataacataaaattaaatcaaatccccctattcccttgatcaattttaatcacagcaattttattaattttatcctta |
| bedd25d69d4bdb2bbf23d3e081e038cf2b873f1c | Hemiptera | Cicadellidae | Sorhoanus | Sorhoanus pascuellus | not found | not found | no | Norway, Germany | Canada, United States of America, Norway, Germany, Netherlands | Possible synonym to Arthaldeus pascuella which is present in Denmark? | cctatctagaaatattgcacatgcaggagcaagagttgatatgtcaatcttttcccttcatttagctgggatctcctctatccttggggcagttaacttcatttcaacagtaataaatatgcgacctacaggaataacactagaccgaaccccactatttgtatgatctgtgttaatcactgccgtcctacttctattgtcttta |
| bac5912832bc40317d7b0014d2cb9b3c5e9d6c9a | Hemiptera | Delphacidae | Nothodelphax | Nothodelphax consimilis | not found | not found | no | no | Canada, United States of America |  | tctgtcaagaatcacttcacattcagggccctcagttgatctaacaatcttttcccttcacatcgctggtgttagttcaattataggagcaattaatttcatctcaacaattatcaatatgcattctaaaactatttcaatagaaaatttacccttattctgttgatcagttttaattacagctattctactactcctttcccta |
| d33fdb3f9a8e4ec299eeba6e6b908762c17e4cb0 | Hemiptera | Miridae | Monalocoris | Monalocoris amamianus | not found | not found | no | no | South Korea |  | tttatcacataatatttcccataatggggcatcagtagatctagcaattttttccttacacttagcaggtatttcatcaatcttaggagcagtaaattttatttcaacaatcatcaatatacgagcaataggaataacagctgatcgtttaccattatttgtgtgatccgtaggaatcacggcattattattattattatcatta |
| 0269283a9458e3f6185a7d6878a4996e886b0c16 | Hymenoptera | Braconidae | Apanteles | Apanteles lacteicolor | no | Germany | not found | not found | Found in 21 countries: <https://fauna-eu.org/cdm_dataportal/taxon/d27a2335-0f4d-48c1-bc33-3277c9eef4e1> |  | attatcattaattttaggacatggaggtatatctgttgatttgggaattttttctttacatttagctggtgcttcatcaattataggggctgttaattttattaccaccattttaattatacgaacaaatttgtttttaatagataaaatatctttattttcttgatctgtttttattacagctattttattattattatcttta |
| 4255f397ee18c4b510cb6961b39b83a1c93fcd7e | Hymenoptera | Braconidae | Ascogaster | Ascogaster provancheri | not found | not found | no | no | Canada, United States of America | Parasitoid on Acleris gloverana and  Acleris variana | tttatctttaattattggtcatggtggtatttcagtagatttaagaattttttctttacatttagctgggatatcttcaattataggtgcaattaattttattgttactattataaatatatgatttggtttaaaatatatagataaaattagattatttacttgatcagtaataattacagcaattttattattattatctttg |
| 1716df57b3c70a71b3bbb5244529725a8a4e266d | Hymenoptera | Braconidae | Chelonus | Chelonus andrievskii | not found | not found | no | Germany | Canada, Turkey, Israel, Germany, Bulgaria | Parasitoid on superfamilies Tortricoidea and Pyraloidea (genus Chelonus) | attgtctttattaataggtcatagaggtgtatcagttgatataagaattttttctttacatttggctgggatatcatcaattataggttcaattaattttattgttactattataaatacatgaataaaatttagttttatagataaatttcctttatttgtttgatcagtttttattacaactattttattgttattgtcttta |
| 695441a8b5e266a90155befb58759639c859642a | Hymenoptera | Braconidae | Cotesia | Cotesia rubecula | no | Germany | no | no | Canada, Australia, New Zealand, Spain, France, Netherlands, in 19 countries: <https://fauna-eu.org/cdm_dataportal/taxon/3a70fd0b-1c49-4bc7-a003-aa09b5da3f3d> | Parasitoid on Pieris rapae | tttatcattaattttaggtcatggtggaatatctgtagatttaggaattttttctttacatttagctggagcatcttctattataggtgctgtaaattttattactactattataaatatacgttcaaatttatttaatatagataaaatatctttattttcttgatctgtatttattactgcaattttattattattatcttta |
| da42908e9e4d66624c0e134ed473c845cc54b7d1 | Hymenoptera | Braconidae | Cotesia | Cotesia vanessae | no | Germany | no | no | In 19 countries: <https://fauna-eu.org/cdm_dataportal/taxon/49568356-3a6a-4e97-a9ff-e13486cb4a1e> | Parasitoid on Noctuidae and Nymphalidae | tttatctttaattttaggacatagtggtttatctgttgatttaagaattttttctttacatttagctggtatatcttcaattataggagcagttaattttattacaacaattttgaatatacgttctggtatatttaatatagataaaatatctttattttcttgatcagtatttattactgcgattttattattattatcttta |
| db4b6d3a6d99b9ed67010be5d34397551fd43cb2 | Hymenoptera | Braconidae | Diaeretellus | Diaeretellus svalbardicum | not found | not found | no | Norway | Norway, Canada | Parasitoid on aphids | cttatctttaactttaggacatagaggtgtggcggtagattttgctattttttcactacatttagcaggtatttcttcaattataggagctattaattttattagtactatttttaatatacgatcatataatattaaaatagatcaaatttcattattaatttgatcagttttaattactgctattttattattattatcatta |
| fd62d3b9b5752d768710e3d5424becf68452de13 | Hymenoptera | Braconidae | Distatrix | Distatrix formosus | not found | not found | no | no | Czechia |  | attatctttaattattggtcatagaggtatatcagttgacataagaatcttttctcttcatttagcaggtgcatcttcaattataggagcaattaattttatttcaacaatttttaatatacgtacttttttttttgaaatagataaaatttcattattttcttgatctgtattaattaccacaattcttttacttttatcatta |
| 8b595436e5845508acf3d1cb1c139a981432e59d | Hymenoptera | Braconidae | Dolichogenidea | Dolichogenidea phaloniae | not found | not found | no | no | United States of America, Canada, United Kingdom, Russia | Parasitoid on Zeuzera pyrina | tttatcattaattttaggacatggtggtatatcagttgatttaggtattttttcattacatttagctggtgcttcatcaattataggtgctgttaattttattacaacaattttaaatatacgaacaaatttatttataatagataaaatatctttattttcttgatcagtttttattactgcaattttattattattatcatta |
| bb3d0f50388e93b2b61a42586c4a249198555d96 | Hymenoptera | Braconidae | Opius | Opius ocreatus | not found | not found | no | Germany | Germany, Turkey, Slovakia |  | attatcatctatggttggtcatggtggtttatcagttgatttagctattttttctttacatttagctggtgtttcttcaatcataggagctattaattttattacaactatttttaatataaatttttatataattaaattagatcaattaagtttattaatttgatcaattttaattacggcaattctattattattatcatta |
| c19882fbf13eced9b80233949ed1189fbb37a851 | Hymenoptera | Braconidae | Protapanteles | Protapanteles porthetriae | no | Germany | not found | not found | found in 23 countries: <https://fauna-eu.org/cdm_dataportal/taxon/734c06c3-edcc-4ac1-8be7-6c11512236df> | Parasitoid on Lepidoptera | tttatcattaattttaggtcatagaggaatatcagttgatatgggaattttttctttgcatttagctggtgcttcttcaattataggtgcagtaaattttattactacaattttaaatatacgaacaaatttatttaaaatagataaaatatctttattttcttgatctgtatttattactgctattttattattattatcttta |
| 6d06367c95f0bd114acb8edbedde4f1370840c9b | Hymenoptera | Braconidae | Remaudierea | Remaudierea plocamaphidis | not found | not found | not found | not found | Czechia, Canada |  | tttatcattaactttaggtcatagaggggttgcagtagattttgctattttttctttacatttagctgggatctcatcaattataggagcaattaattttattagtactatttttaatatacgttcttataatattaaaatagatcaaatttctttattagtttgatcagttttaattacagctgttttattattattgtcttta |
| 63e137c75dea6f47b4bfab5c5f2305b38fccb2ec | Psocodea | Liposcelididae | Liposcelis | Liposcelis rufa | no | no | no | no | Found in 15 countries: <https://fauna-eu.org/cdm_dataportal/taxon/13f0b344-cf15-480b-997a-4941ae0c8682> |  | ctatcattatattcagcccatcccagtgaaagtgtagacttagcaattttttctcttcacttagctggagcgagatcgattttaggagctattaattttattactacattttttaatctatgagtttttaaaaataaaatagaattgagaagtttatattcttggtctgtttcaatcacagcagttttacttttactttcttta |
