## Supplementary_Material for "Detecting flying insects using mega-nets and meta-barcoding"

### Car net design

Our car net was made of 100 % polyester bunting with a PVC mesh with side stiffeners in the bottom enabling a steady airflow and expanding the opening of the sampling bag when the car is in motion. The opening of the car net is secured by aluminium poles custom manufactured to increase resistance to wind pressure and to blunt the ends of the tent poles to decrease wear on the net. Metal guy line adjusters enabled adjustment to car length and allowed the net to be used on most car types. The end of the main net is a detachable (velcro) sampling bag with a closing guy line adjuster sewn into each sampling bag (Fig S1). A 5 cm wide magnetic band was sewn into the bottom of the net opening to prevent the net from lifting from the roof of the car and prevent insects “escaping” under the net.

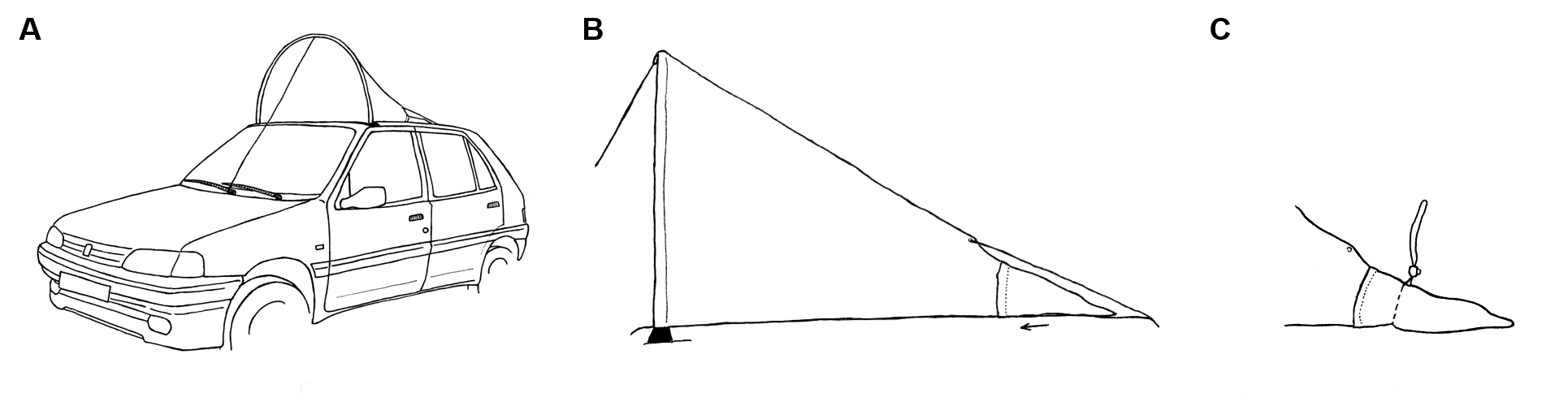

Figure S1 Drawing of the car net used to sample insects. A) the car net mounted on the car with a magnetic strip at the bottom of the front opening to secure the opening to the roof of the car and adjustable straps going into the car through the passenger doors. B) Side view of the car net with adjustable straps at the front and back, and C) the detachable sampling back at the back end of the large net, with adjustable pull string to close the back and secure the collected insects. Drawings by Marie Rubæk Holm.

### Laboratory methods

The full laboratory protocol can be accessed at [dx.doi.org/10.17504/protocols.io.bmunk6ve](https://dx.doi.org/10.17504/protocols.io.bmunk6ve).

#### Insect abundance: order level

Bulk insect sample handling and DNA extraction were carried out in dedicated pre-PCR extraction laboratories. In a laminar flow hood, the content of the sample bag was poured into a sterilized glass tub with a laminated sheet of millimeter paper under the tub. Insets still attached to the sampling bag were removed with a squeeze bottle containing 96 % ethanol and sterile forceps. Insect abundance was assessed by counting individuals and assigning them by morphology to order level with a stereomicroscope. Excess ethanol was removed by pouring the sample through a cleaned vacuum filtration device with one-use filters (Whatman™ grade 4 and 4V). Filter blanks were included in preliminary analysis to ensure the filtration method did not introduce cross contamination between samples. Leftover ethanol was evaporated at 50˚C in an oven prior to DNA extraction and an extraction blank was included for every 8-12 samples.

#### DNA metabarcoding of bulk insect samples

##### DNA extraction and qPCR

The dried insects were extracted with a non-destructive DNA buffer (Nielsen et al., 2019) modified from (Gilbert et al., 2007) which leave the exterior of most insect taxa intact but dissolves the interior of the insects, including gut content. The volume of dried insects in the tube was used to calculate the amount of digestion buffer added to each sample so final DNA concentrations were as even as possible between samples. The samples were lysed at 50°C overnight mounted on a rotator in an oven and centrifuged in a SL 40 centrifuge (Thermo Fischer Scientific) for 2 min. at 2000 x g to pellet insects. Digest was removed and the insects were washed with 96 % ethanol to stop the digestion. Ethanol used for washing was removed, and new 96 % ethanol was added to preserve the insects as bulk sample vouchers within the Natural History Museum of Denmark’s entomological collection. DNA extracts were stored at -20°C. Automated purification was carried out for an aliquot of 230 µl digest from each sample with magnetic-particle purification on a QIAsymphony SP robot (Qiagen). 23 samples and one extraction blank was included in each purification run and the DSP DNA Mini Kit (Qiagen) and the Tissue_LC_200_V7_DSP protocol (Qiagen) was used for all samples. DNA concentration of the purified samples were quantified with QuBit® 3.0 Fluorometer (Thermo Fisher Scientific) and an aliquot of purified DNA was normalized to 1 ng/µl DNA input prior to PCR. Prior to PCR, a dilution series was run on a Stratagene MX3005P qPCR machine (Agilent Technologies) to determine the optimal DNA input and number of cycles for PCR.

##### PCR amplification

A universal insect primer was used for DNA metabarcoding targeting CO1, a mitochondrial DNA (mtDNA) gene frequently used in arthropod studies. The CO1 primer set used was fwhF2 + fwhR2n (~205 bp) (Vamos et al., 2017). The primer set was tagged with 96 unique forward and reverse tags, which were composed of six nucleotides including one to three random bases to add complexity. PCR reactions of ~25 µl were run with Thermo Fisher Scientific AmpliTaqGOLD Master Mix (2.5 µl 10X PCR GOLD buffer, 2.5 µl GOLD MgCl2 (25 mM), 0.2 µl dNTPs (25m M), 0.2 µl AmpliTaqGOLD polymerase (5U/µl), 1 µl BSA (20 mg/mL), 11.6 AccuGENE molecular grade water, and 1.5 µL forward and 1.5 µl reverse primer (10 µM) with 4 μl of diluted DNA aliquots. The thermal cycling profile was 95°C for 5min followed by 34 cycles of 95°C for 30 s, 58°C for 30 s, 72°C for 45 s, with a final extension time of 72°C for 10 min. All PCRs were run with two PCR and DNA extraction negatives to determine the potential presence of contamination. The amplified PCR products (5 μl) were visualised on 2 % agarose gel stained with GelRed^TM^ by electrophoresis. To ensure all samples were equally proportioned, the PCR products were pooled in different amounts depending on the intensity of the bands visualized on the gel. Scoring based on the gel visualization was evaluated with QuBit^®^ for a subset of PCR products. Samples with low-intensity bands (7.5 μl aliquots), medium intensity bands (5 μl aliquot) and high-intensity bands (2.5 μl aliquot) were pooled together prior to library build. Tags were used only once in each pool to eliminate sample mismatches due to tag jumps. Pooled samples were purified with MinElute PCR Purification Kit as preparation for library build.

##### Library building & next-generation-sequencing

Two libraries of 96 samples each were built using the TruSeq DNA PCR-Free Library Preparation Kit (Illumina) with TruSeq DNA UD Indexes (Illumina). The libraries were purified with MagBio LabLife HighPrep PCR beads to remove primer dimers. To determine the length of products and concentrations for pooling, the libraries were run on an Agilent Technologies 2100 Bioanalyzer. Libraries were pooled in equimolar concentrations and sequenced on an Illumina NovaSeq 6000 (150 bp paired-end) sequencing platform (Illumina Inc., San Diego, CA, USA) with a spike-in of 4 % PhiX to increase complexity.

##### Bioinformatic analysis & taxonomic assignment

Sequencing libraries were demultiplexed using cutadapt (version 1.11) (Martin, 2011) and processed with the DADA2 pipeline (version 1.8) (Callahan et al., 2016) in RStudio (version 3.6.5) to detect erroneous sequences generated during PCR amplification and sequencing. DADA2 analyses were performed on each FASTQ file separately and forward and reverse reads were merged with a minimum overlap of 5 bp. Likely chimeras were removed with the DADA2 function removeBimeraDenovo. BLASTn was used to make a LULU match list from the blast database and processed with the post clustering algorithm LULU (Frøslev et al., 2017) to flag and remove erroneous OTUs. Sequences with a length ≥200 bp were taxonomically assigned. The Global Biodiversity Information (GBIF) sequence ID tool (https://www.gbif.org/tools/sequence-id) was used to assign taxonomy to CO1 sequences. The sequence ID tool queries a 99% clustered version of the International Barcode of Life (iBOL) public data sequence database and uses GBIFs backbone taxonomy to assign names. Observations that matched class Insecta were used for the primary analysis, since we assume unique sequences represent true diversity. ASVs that matched class Insecta and had a query coverage of ≥80% and identity match ≥99% to the reference were included for proportional visualisation, as an evaluation of the reference database suitability for diversity estimates.

### Results

#### Sequence data results

DADA2 processing yielded 32050 ASVs which were reduced to 16182 ASVs (50.5%) in 1058 samples by applying the LULU algorithm which flags and, in this case deletes, potentially artificatial ASVs. Of those, 13799 ASVs (85.3%) were retained after filtering read length to >200 bp. Of the 13799 ASVs, 8172 matched the class Insecta and were retained for further analysis. Filtration by a ≥99% identity match to the reference sequence yielded 5698 (69.7%; these ASVs corresponds to 17.8% of the ASV output from DADA2). The 5698 ASVs corresponded to 1829 uniquely named species because some ASVs did not have species level matches in the database. Of the 1058 samples initially sequenced, we retained the 365 samples from 2018 for analysis.

#### Species accumulation curve

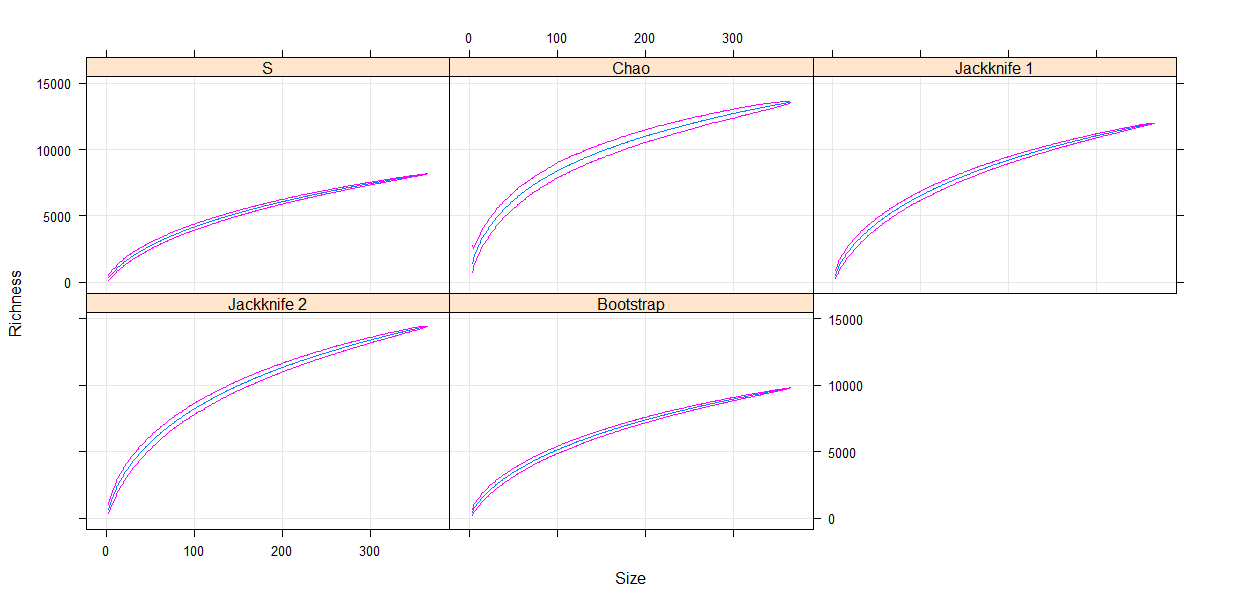

Fig S2 Output from the poolaccum function with different estimators calculation of richness for each sample (cumulative) (permutations = 1000). S is the observed ASV richness.
